# Size-tunable ICG-based contrast agent platform for targeted near-infrared photoacoustic imaging

**DOI:** 10.1101/2022.09.01.506234

**Authors:** Shrishti Singh, Giovanni Giammanco, Chih-Hsiang Hu, Joshua Bush, Leandro Soto Cordova, Dylan J Lawrence, Jeffrey L Moran, Parag V Chitnis, Remi Veneziano

## Abstract

Near-infrared photoacoustic imaging (NIR-PAI) combines the advantages of optical and ultrasound imaging to provide anatomical and functional information of tissues with high resolution. Although NIR-PAI is promising, its wide application is hindered by the limited availability of NIR contrast agents. J-aggregates (JA) made of indocyanine green dye (ICG) represents an attractive class of biocompatible contrast agents for PAI. Here, we present a facile synthesis method that combines ICG and ICG-azide dyes for producing contrast agent with tunable size down to 230 nm and direct functionalization with targeting moieties. The ICG-JA platform has a detectable PA signal *in vitro* that is two times stronger than whole blood and high photostability. The targeting ability of ICG-JA was measured *in vitro* using HeLa cells. The ICG-JA platform was then injected into mice and *in vivo* NIR-PAI showed enhanced visualization of liver and spleen for 90 minutes post-injection with a contrast-to-noise ratio of 2.42.

## 1. Introduction

Photoacoustic imaging (PAI) [1–3] is a relatively low-cost non-invasive biomedical imaging modality, which combines the advantages of optical imaging (e.g., high contrast and molecular specificity) and ultrasound imaging (e.g., deep penetration in tissue). Near-infrared I (NIR-I) illumination (620-950 nm) can be used with PAI to reduce photon scattering in biological tissues and improve imaging depth [4]. To this end, several exogenous contrast agents (CA) such as organic particles, metallic nanoparticles, [5,6] or polymer-based nanomaterials [7,8] have been designed for NIR-PAI to enhance the image contrast. Hence, NIR-PAI has found use in various pre-clinical applications, including imaging tumor lesions, [9] detecting metastatic lymph nodes, [10,11] delineation of tumor margins, [12,13] identification of pathological tissues, [14] and measurement of blood oxygenation level [15]. However, even with the current catalogue of available CAs, NIR-PAI still requires improved CAs that combine critical features such as stability *in vivo*, solubility, biocompatibility, photostability and targeting capabilities [16,17]. In addition, there is also a need to develop robust and simple methods that reproducibly synthesize different sized CAs depending on the biological system and applications [18,19]. Thus, it is essential to engineer a stable and biocompatible CA platform that can enable prescribed size tuning and easy functionalization with targeting moieties [16].

One of the main approaches taken for producing CA of a specific size and functionalization is to produce aggregates of dyes [20]. Indocyanine green (ICG) is an attractive candidate for this purpose since it is a biocompatible FDA-approved NIR dye currently used for PAI [9]. Monomeric ICG dye has variation in its optical properties depending on its surrounding environment [21], has low photostability and its targeting chemistry is difficult [22]. On incubation in aqueous solutions at temperatures above 5°C [23], ICG can assemble to form J-aggregates (JA) [24,25]. J-aggregates are stable head-tail arrangements of monomeric dye held together by non-covalent interactions with a strong red (bathochromic) shift of the absorbance peak coupled with higher absorbance [26,27]. Compared to monomeric ICG, ICG-JA have advantages like quenched fluorescence [28], a red-shifted absorption peak at a wavelength of 895 nm[29], stronger PAI signal, [30] and greater photothermal stability [31]. However, the formation of ICG-JA is a process of self-assembly of monomeric ICG, which often yields micron-sized, polydisperse aggregates. Therefore, most current methods of synthesizing monodisperse, nanometer-sized ICG-JA rely on syringe filtration [28,29] or encapsulation in nanocarriers [24,32,33]. The addition of targeting moieties also requires encapsulation in nanocarriers. These factors limit the flexibility of ICG-JA with respect to size, targeting and photoacoustic properties.

Hence, in this study, we report the scalable preparation of azide-modified ICG J-aggregates (JAAZ) that have controllable mean sizes ranging from 230 to 1200 nm, are amenable to direct functionalization, with NIR absorption, and a strong PA signal. As a proof of principle, we functionalize RGD peptide as a model of targeting moiety to enable targeting of cells that overexpress the integrin receptor such as HeLa cells. We determined the intensity of the PA signal in an agarose phantom and upon binding to HeLa cells. We further evaluated the biodistribution, PA properties and capability to visualize blood-rich organs of RGD functionalized JAAZ with preliminary experiments *in vivo* using tomographic PAI.

## 2. Results and Discussion

### 2.1. Synthesis, characterization and stability of azide-modified J-aggregates

To synthesize azide-modified ICG-JA (JAAZ), we first incubated different ratios of ICG and ICG-azide dye in water using a procedure previously described for ICG only (Figure 1A) [29]. After incubation for 20 and 40 hours at 60°C in water, we observed the typical JA peak at 895 nm for 1:10 ICG-azide:ICG solution but not for 100% ICG-azide dye or for ICG-azide:ICG molar ratios of 3:4, 1:2 and. 1:4 (Figure S1). However, for the JA formed using the 1:10 ratio, the Fourier-transform Infrared (FT-IR) spectrum did not show a peak between 2141-2100 cm^-1^ (Figure S2), which is a characteristic range for the azide group due to the strong asymmetric N=N=N stretching, [34] indicating that the formed JA did not contain ICG-azide but only ICG. To promote the incorporation of ICG-azide into the JA, we modified our initial protocol to include cations, which are known to influence aggregate formation [35]. Indeed, monovalent metal cations like Na^+^ and K^+^ are known to facilitate J-aggregate formation of anionic cyanine dyes such as ICG and pseudoisocyanine (PIC) due to ionic interactions between cations and the dye molecules that limit electrostatic repulsion between dyes [36,37]. Thus, we tested different concentrations of potassium chloride (KCl) (1, 5, 10 and 20 mM) with 1:10 ICG-azide:ICG molar ratio solutions. Additionally, we also tested formation of JAAZ particles with different molar ratios of ICG-azide to ICG (1:10, 1:5 and 1:3) using 20 mM KCl. After 20 hours of incubation at 60°C, all conditions showed the presence of a red-shifted absorption peak at 895 nm and the disappearance of both the monomer and the dimer peaks (780 and 715 nm, respectively) (Figure 1B, Figure S3 and S4A). FT-IR spectra of all synthesized JAAZ particles with KCl show a peak at 2100 cm^-1^ due to incorporated azide groups, which is absent from the control ICG-JA (Figures S4B-D). We further studied temperature as an experimental parameter effecting the formation of JAAZ particles. The formation of JAAZ particles was quantified by the intensity of the 895 nm absorption peak (Figure S5) at 50 °C, 60 °C and 70 °C. At 50 °C, the intensity of the 895 nm was lower than 60 and 70 °C at 20 hours of incubation. At 70 °C, the intensity decreased after 14 hours indicating the slight decomposition of ICG dye. Hence, incubation at 60 °C for 20 hours was appropriate for JAAZ formation.

**Figure 1.**
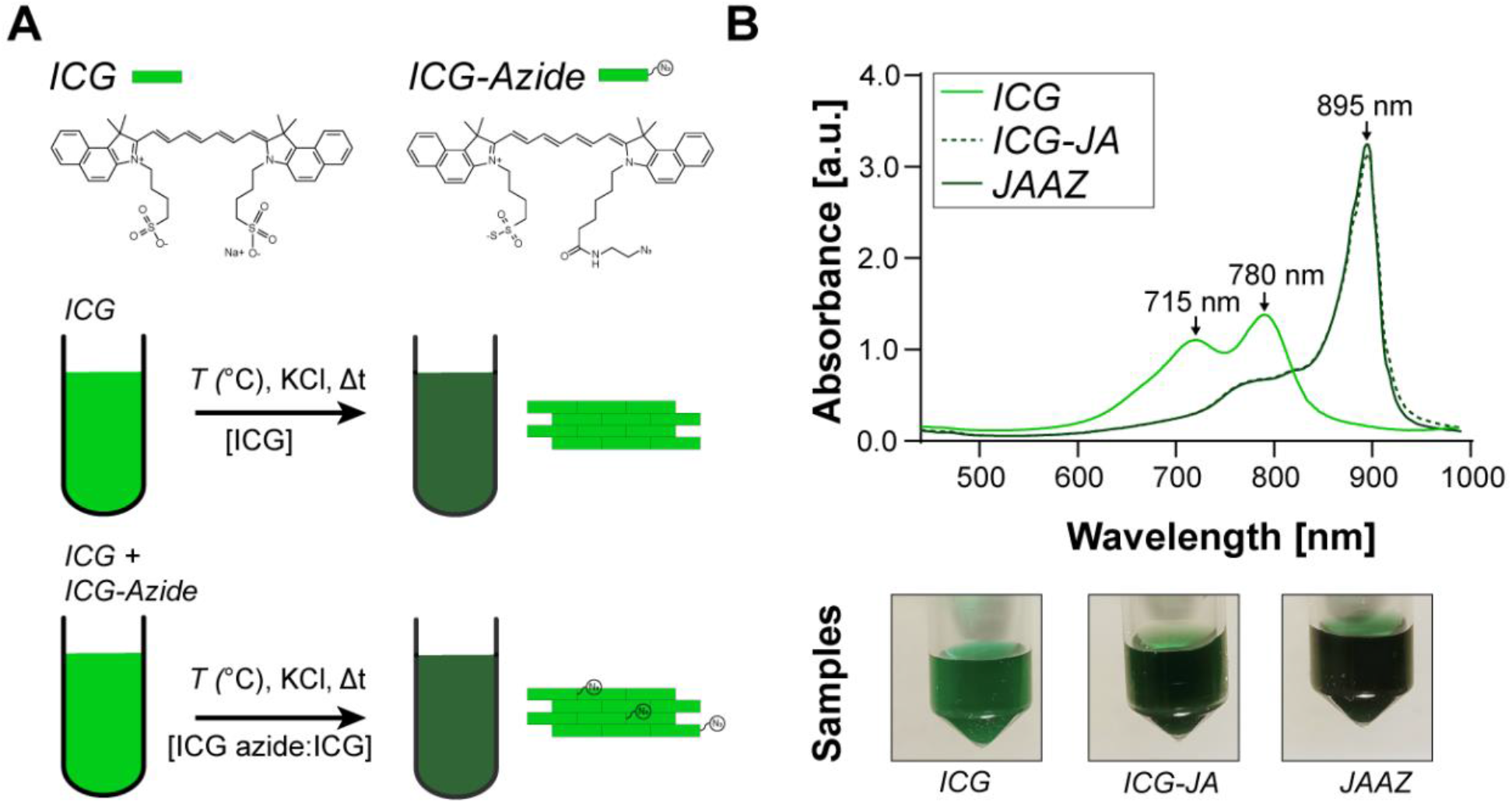
Synthesis of ICG J-aggregates (ICG-JA) and azide-modified ICG J-aggregates (JAAZ) particles. (A) Schematic of the procedure used to synthesize ICG-JA and JAAZ particles. (B) (top) Representative absorbance spectra of ICG-JA and JAAZ particles at 50 μM of total dye concentration. A change in the color of the solution is observable by the naked eyes between monomeric ICG solution and the ICG-JA and JAAZ particle solutions due to the shift toward longer wavelength.

Before further functionalization with targeting moieties, excess free dye and residual salt were efficiently removed from the prepared ICG-JA and JAAZ using spin column filtration (Figure S6A). There was no fluorescence emission at 800 nm for both the ICG-JA and JAAZ particles as previously described in literature [28] (Figure S6B), which is an important characteristic since it insures maximum conversion of the absorbed photons to acoustic signal. Therefore, different experimental parameters like KCl concentration, temperature and ICG-azide to ICG molar ratio facilitated the formation of J-aggregates with incorporated azide groups.

After confirming formation of JAAZ particles, we examined the presence and accessibility of the azide group using measurements of size and surface charge. We first measured the surface zeta (ζ) potential of both ICG-JA and JAAZ, knowing that the difference in charge between ICG-azide dye and regular ICG dye should modify the overall surface charge of the JAAZ particles if ICG-azide dyes were to be displayed on the surface. The average ζ potential measured was −50.32 mV and - 60.11 mV for the JAAZ and the ICG-JA particles, respectively (Figure 2A). The less-negative zeta potential of JAAZ particles indicates the presence of azide groups on the surface of the JAAZ particles. Moreover, the narrow surface charge distribution confirms the presence of a single J-aggregate population (Figure S7). Using dynamic light scattering (DLS), we measured the average hydrodynamic diameter of ICG-JA and JAAZ particles as 1.17±0.26 and 1.34±0.18 μm, respectively (Figure 2B). The slightly larger size of JAAZ particles could be attributed to a less densely-packed or organized arrangement of ICG molecules due to the presence of azide groups. Scanning electron microscopy (SEM) reveal that JAAZ particles have a uniform morphology with an average diameter of 1.1 μm, in good agreement with the DLS data (Figures 2C and S8A). Energy-dispersive X-ray (EDX) spectroscopy confirmed the elemental composition of JAAZ particles (Figure S8B).

**Figure 2.**
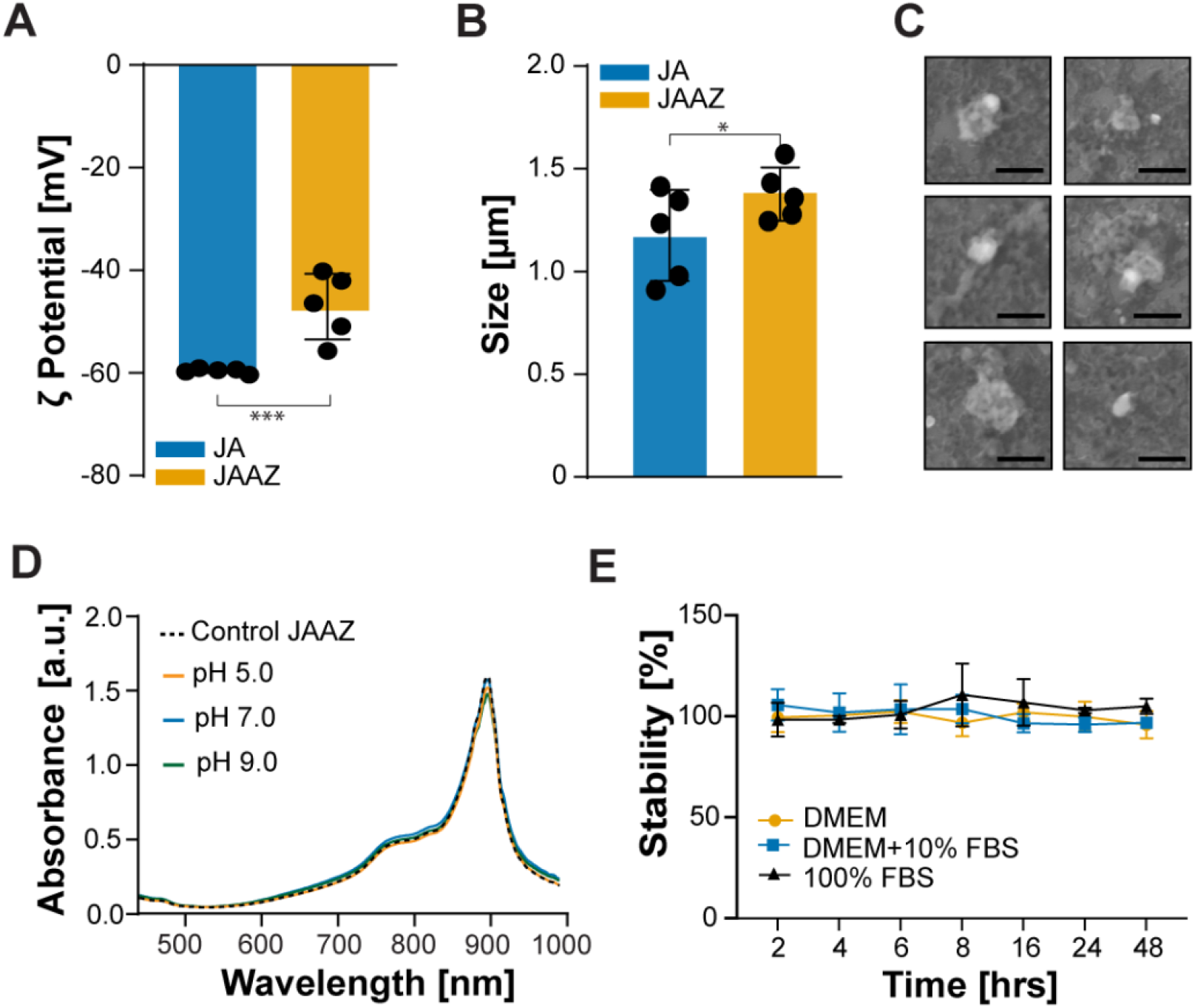
Characterization of JAAZ particles prepared in 20 mM KCl at 1 mM dye concentration. (A) Zeta potential measurements of ICG-JA and JAAZ particles (B) Dynamic light scattering measurements of ICG-JA and JAAZ particles (C) SEM images of JAAZ particles (Scale bar: 2 μm, the complete SEM image is provided in Figure S8). (D) UV-Vis-NIR absorbance of JAAZ particles incubated in solutions of pH 5 (red), 7 (blue), and 9 (green) at 37 °C for 126 hours. (E) Percentage stability of JAAZ particles in DMEM (blue), DMEM+10%FBS (orange) and 100% FBS (black). Stability over time was quantified as difference in the intensity of the 895 nm absorption peak when compared to control at 0 minutes. (n= 5 for A and B, n= 3 for D and E, all results are mean±SD with p value calculated using unpaired t-test; *and ***=p<0.05, all P values provided in Table S3).

In order to determine the stability of the physical properties of JAAZ particles, we lyophilized and resuspended the JAAZ particles in water. There was no significant change in the absorption spectra, size and ζ potential, suggesting that JAAZ particles can be stably stored in lyophilized form at room temperature protected from light (Figure S9 and Table S1). Moreover, they are easily dispersed in different media, such as 1X phosphate buffered saline (PBS), Dulbecco’s Modified Eagle Medium (DMEM), DMEM+10% fetal bovine serum and 100% fetal bovine serum (FBS, mimicking biological conditions). JAAZ particles were stable in 1X PBS at 37°C and different pH (5, 7 and 9) for up to 126 hours (Figure 2D). They were also stable in DMEM, DMEM+10% FBS and 100% FBS for 48 hours as observed by the lack of intensity difference in the 895 nm peak (Figure 2E), indicating JAAZ particles were stable in *in-vitro* physiological conditions.

Since J-aggregates are an assembly of dye molecules, it was important to determine the concentration of dye within the cluster before functionalization. Hence, we disassembled the J-aggregates into free dye using Triton X-100 to determine the equivalent total dye concentration and compare it from one sample to another. A similar concentration of dye was determined using a standard curve (Figure S10) for both ICG-JA and JAAZ particles, indicating that the efficiency of azide-modified J-aggregate formation is similar to ICG-JA.

### 2.2. Functionalization of the azide-modified J-aggregates

To enable further functionalization of JAAZ with streptavidin, we performed copper-free click chemistry to graft DBCO-PEG-biotin linker onto the azide groups available on their surface to generate biotinylated JAAZ particles (Bio-JAAZ). After removal of the excess of DBCO-biotin linker, we incubated the particles with streptavidin to prepare Strep-JAAZ. Biotinylated RGD was then attached to available streptavidin to create RGD-JAAZ, as shown in Figure 3A. A summary of the concentration of biomolecules used to functionalize JAAZ to create Bio-JAAZ, Strep-JAAZ and RGD-JAAZ is provided in Table S2. Conjugation of the aggregates with biotin, streptavidin and RGD did not affect the aggregates stability as demonstrated with the absorbance spectra of Bio-JAAZ, Strep-JAAZ and RGD-JAAZ showing the typical J-aggregate absorption peak at 895 nm (Figure 3B). Bio-JAAZ particles exhibit an average ζ potential of −54.1 mV, a non-significant change compared to JAAZ particles. In contrast, the attachment of streptavidin changed the average ζ potential of the particles to −15.8 mV, making them significantly less charged than JAAZ (Figure 3C) as expected given the charge of streptavidin [38]. DLS measurements show the size of Bio-JAAZ, Strep-JAAZ and RGD-JAAZ particles were 1.26±0.34 μm, 1.51±0.35 μm and 1.72±0.36 μm, respectively (Figure S11A). Attachment of streptavidin caused no statistically significant increase in the size of JAAZ particles, confirming the absence of aggregation of Strep-JAAZ particles. Both Strep-JAAZ and RGD-JAAZ particles have a PDI of 0.26 when compared to JAAZ particles with a PDI of 0.37 (Figure S11B). SEM images and EDX analysis (Figure 3D) show the average size of Strep-JAAZ particles as 917.11 μm and confirm the coating of streptavidin due to a significant increase in weight percentage of elements such as carbon, nitrogen and oxygen compared to uncoated JAAZ. To quantify the availability of biotin groups, we attached streptavidin to JAAZ particles with different ratios (1:10, 1:5 and 1:3 ICG-azide:ICG molar ratio) and measured the concentration of streptavidin attached using a bicinchoninic acid (BCA) assay. The amount of streptavidin attached to 1:10, 1:5 and 1:3 JAAZ particles were 166.2, 623.1 and 510.6 μg/ml per 50 μM of equivalent free dye, respectively. JAAZ particles with no biotin groups demonstrate some non-specific binding of streptavidin but with ~70% less streptavidin present when compared to the 1:10 molar ratio Bio-JAAZ particles. For all the ICG-azide:ICG ratios tested for conjugation with streptavidin, we did not observe change in the absorbance peak and the zeta potential (Figures S12A and S12B). These results indicate that a higher degree of biomolecule functionalization can be achieved by varying the molar ratio of ICG-azide:ICG without affecting the stability of the aggregates. Fluorescence spectroscopy was used to confirm the drastic reduction of fluorescence emission for the RGD-JAAZ particles as observed with JAAZ, indicating that addition of biomolecules had no effect on the spectroscopic properties of the aggregates and does not induce their degradation (Figure S6).

**Figure 3.**
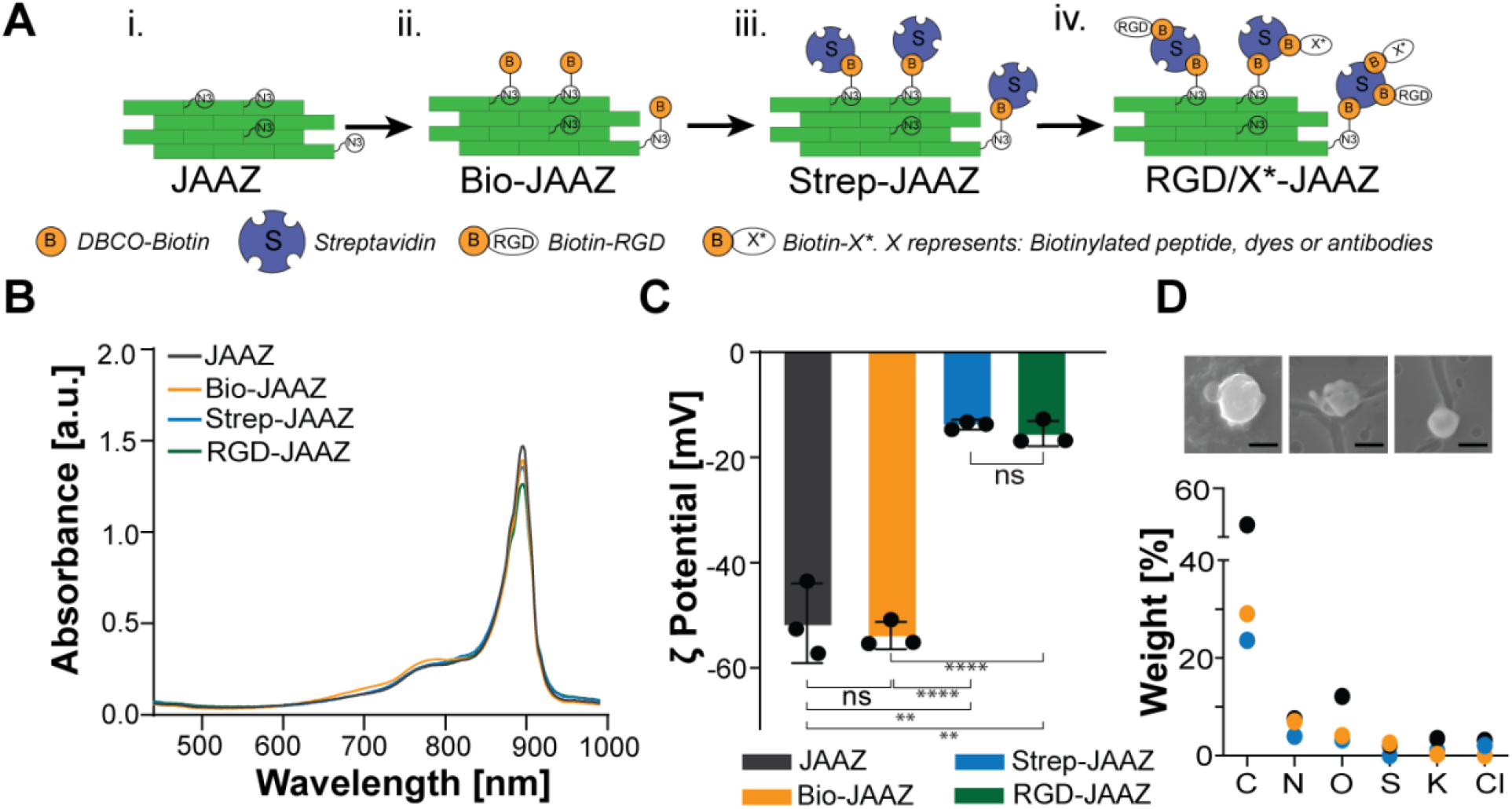
Functionalization of JAAZ particles with biomolecules. (A) Schematic of the JAAZ particles coating with streptavidin. i. JAAZ particles are assembled and purified. ii. Biotin-PEG-DBCO is attached to the azide groups via click chemistry to form Bio-JAAZ. iii. After purification of the Bio-JAAZ, streptavidin is attached to the biotin group to form Strep-JAAZ. iv. The introduced streptavidin can then be used to attach any biotinylated targeting moieties such as peptides (e.g, RGD-biotin). (B) Absorbance of functionalized JAAZ samples. (C) Zeta potential measurements of functionalized JAAZ particles. (D) SEM images and EDX analysis of streptavidin coated JAAZ particles (Scale bar: 1 μm). (n=3 for B and C, data presented in C represented as the mean±SD, p-values are calculated using one-way ANOVA, **=p<0.01 ****= p <0.0001; all P values provided in Table S4)

We observed that modification of JAAZ particles with a soluble protein like streptavidin increased their solubility, preventing sedimentation. To test this, JAAZ and Strep-JAAZ particles were left to sediment at room temperature on a benchtop. No sedimentation was observed for up to 8 hours with Strep-JAAZ particles, whereas a visible pellet was formed for the JAAZ particles. At 24 hours, Strep-JAAZ particles settled down at the bottom of the centrifuge tube. However, upon slight mixing, they returned into solution, whereas a visible pellet was observed for JAAZ particles at 24 hours, which required vortex mixing to be properly dispersed (Figure S13A and S13B). The results presented in this section demonstrate our capability to conjugate biomolecules to the

### 2.3. Synthesis and characterization of nanosized azide-modified J-aggregates

Because the micrometer size of JAAZ particles might restrict their use in applications that require contrast agents to pass through biological barriers such as the blood brain barrier, we adapted our synthesis protocol to enable control over the size of JAAZ particles. According to previous literature, [23,35,39] J-aggregate formation is dependent on time, salt and dye concentration; hence, we explored the impact of these three parameters and their combination on the size of these JAAZ particles. Solutions of 250, 500, 750 and 1000 μM total dye with 1:10 ICG-azide:ICG molar ratio were incubated at 60°C for 20 hours under KCl concentrations of 0.1, 1 and 20 mM. At 0.1 mM KCl, the 895 nm absorption peak appeared after incubation for 6 hours for 1000 and 750 μM of total dye. For 500 and 250 μM concentrations, 8 and 14 hours were required for the formation of J-aggregates, respectively (Fig. S14A-D). For 1 and 20 mM KCl, a J-aggregate peak formed after 6 hours for all dye concentrations (Figure S15 and S16). The size of the formed JAAZ particles was measured using DLS for the above-mentioned conditions after 20 hours. At higher total dye concentrations for all salt molarities, the size of the formed JAAZ particles ranged from 700 nm to 1.4 μm. On reducing the initial concentration of total dye, at lower salt molarities of 0.1 mM and 1 mM KCl, nanoscale JAAZ (N-JAAZ) of average size ranging from 230 to 600 nm were formed (Figure S18A-D). Using total dye concentrations of 500 and 250 μM in 0.1 mM KCl, we demonstrated the synthesis of 350 and 225 nm JAAZ nanoparticles, respectively (Figure 4A). These results indicate that varying the concentration of total dye, concentration of KCl and incubation time can be used to tune the size of JAAZ particles. Hence, we were able to synthesize N-JAAZ particles with average size of 225 nm (Figure 4B) and a PDI of 0.20 with 250 μM total dye and 0.1 mM KCl. Using a higher concentration of KCl and total dye (1 mM KCl, 750 μM dye), JAAZ particles of 640 nm were obtained (Figure 4B). JAAZ particles of 1.3 μm, as shown in Figures 2B and 4A, were formed in a solution of 20 mM KCl, 1000 μM total dye. A detailed summary of the size of JAAZ particles formed varying time, KCl and total dye concentrations is given in Figure S18. To demonstrate the difference in sizes between the JAAZ and N-JAAZ, the two samples were loaded onto an agarose gel where the negatively charged JAAZ and N-JAAZ particles were expected to move toward the positive electrode. N-JAAZ particles of ~230 nm moved through the gel and formed a distinct green band after traversing the gel as seen in Figure 4C, whereas the size of JAAZ particles (~1 μm) prevents them from migrating through the gel. The size of N-JAAZ particles was further confirmed by SEM images (Figure 4D) which showed particles with an average diameter of 175 nm. EDX analysis (Figure S19) confirmed the composition of N-JAAZ particles.

**Figure 4.**
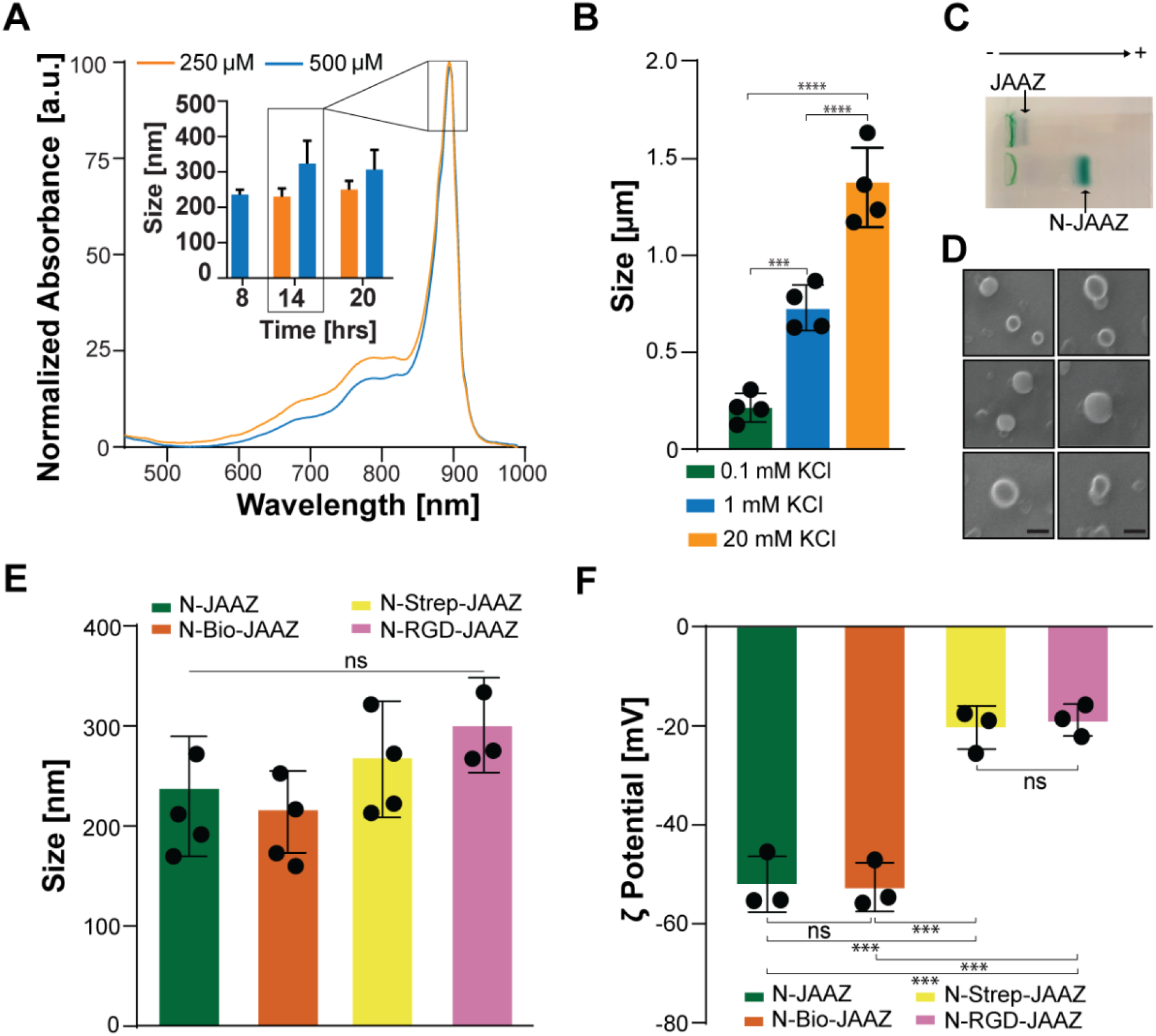
Characterization of N-JAAZ. (A) Absorbance spectrum of N-JAAZ particles formed in 0.1 mM KCl using 1:10 ICG-azide:ICG solutions at 250 and 500 μM of total dye. Representative absorbance is particles formed at 14 hours of incubation at 60 °C. Inset: Size of N-JAAZ after formation for 8, 14 and 20 hours. (B) Size of N-JAAZ and JAAZ particles. The total concentration of dye used for 0.1 mM KCl samples was 250 μM and for 1 mM and 20 mM KCl samples, the dye concentration was 1 mM. (C) Representative agarose gel electrophoresis of JAAZ and N-JAAZ particles. (D) SEM images of N-JAAZ particles prepared in 0.1 mM KCl with 250 μM of dye. (Scale bar: 250 nm, complete image is provided in Figure S20). (E) DLS measurements of functionalized N-JAAZ particles yield average hydrodynamic diameters of 206.6 nm, 256.5 and 313.0 nm, respectively. (F) Zeta potential values of functionalized N-JAAZ particles are −52 mV, −53.1 mV, −20.6 and 18 mV respectively. (n=4 for A, E and F, all data presented as mean±SD, p-values are calculated using one-way ANOVA, ***=p<0.001 and ****=p<0.0001, ns, not significant; all P values are provided in Table S5).

Following the same biomolecule functionalization strategy used for JAAZ particles (Figure 3A), we attached biotin and streptavidin onto the surface of N-JAAZ. After attaching DBCO-Biotin and streptavidin, the average hydrodynamic diameter of N-Bio-JAAZ and N-Strep-JAAZ particles were 210±30 nm and 255±27nm, respectively. After RGD attachment, the average size of N-RGD-JAAZ particles was 313±37 nm (Figure 4E), a statistically non-significant increase compared to N-JAAZ particles. Hence, the synthesized N-JAAZ particles can be functionalized with RGD yielding nanosized and targeted J-aggregates. As in the case of JAAZ particles, attachment of the slightly negatively charged streptavidin to N-JAAZ particles makes their zeta potential more positive. The average ζ potentials of N-JAAZ, N-Bio-JAAZ, N-Strep-JAAZ and N-RGD-JAAZ particles were −52 mV±5.1, −53 mV±5.7, −21 mV±3.3 and −18±2.4 mV respectively (Figure 4F), comparable to those obtained for JAAZ particles. To determine if the reduction in size resulted in changes in the stability of N-JAAZ particles, we performed stability studies in DMEM, DMEM+ 10% FBS and 100% FBS. When incubated in DMEM and DMEM+10% FBS for 24 hours at 37 °C, no significant differences were observed in the intensity of the absorbance peak at 895 nm. However, there was a 20% decrease in stability of N-JAAZ particles in 100% FBS at 24 hours of incubation (Figure S20). In addition, no significant change in absorbance, size or ζ potential were observed for N-JAAZ particles on vacuum drying and resuspension (Figure S9 and Table S1), indicating that a reduced size has no notable influence on the stability of the aggregates. Thus, we have designed an experimental protocol to control the size of JAAZ particles synthesizing stable N-JAAZ particles, which can be modified with biomolecules making targeted nanoscale J-aggregates.

### 2.4. Stability and cytotoxicity of the JAAZ and N-JAZZ particles *in vitro*

We also tested the hemotoxicity of JAAZ, N-JAAZ and RGD-JAAZ particles (Figure S21A). Optical microscopy images of smeared red blood cells stained with all three types of JAAZ particles showed no difference in morphology compared to control blood indicating no adverse reaction of all combinations of JAAZ particles. To assess the cytotoxicity of different JAAZ particles, we performed an MTT assay using HeLa cells (Figure S21B). HeLa cells were incubated with JAAZ, N-JAAZ and RGD-JAAZ at an equivalent dye concentration of 20 μM. With all three samples at high concentration, cells did not show any cytotoxicity after 24 hours of incubation. These data indicate the *in-vitro* safety and efficacy of JAAZ particles.

Before testing the *in vitro* PA properties of JAAZ and N-JAAZ particles, we determined their stability in whole blood (Figure S24). JAAZ particles were ~60% stable when incubated in whole blood at 37°C for 24 hours. N-JAAZ particles were 90% stable degradation at end of the 24-hour incubation. RGD-JAAZ particles were stable for 24 hours, an increased stability when compared to uncoated particles.

### 2.5. Characterization of the photoacoustic properties of JAAZ particles *in vitro*

To quantify their use for PAI applications, we characterized the *in vitro* photoacoustic properties of JAAZ and N-JAAZ along with RGD-JAAZ. Figure 5A shows the photoacoustic setup used, which includes the different J-aggregate samples embedded in agarose and submerged in a water reservoir, illuminated by a pulsed laser. PA signal intensities were recorded for JAAZ, N-JAAZ, and RGD-JAAZ particles at 25, 12 and 5 μM equivalent free dye concentration in 0.5% (w/v) agarose. All PA signals were normalized with respect to 1:100 dilution of India Ink in water. The normalized PA signal magnitude of RGD-JAAZ particles was comparable to the PA magnitude of the non-functionalized particles, demonstrating that attaching a protein like streptavidin does not substantially affect the PA conversion efficiency of the J-aggregates (Figure 5B). To ascertain whether the size of the JAAZ particles have any influence on the PA signal intensity, the PA signal of different sized JAAZ particles (960 nm-250 nm) at the same absorbance intensity at 895 nm was recorded (Fig S22). The PA signal intensity for the different sized JAAZ particles had significant statistical difference, but the overall PA signal intensity for all JAAZ particles was in the same range, signifying that the statistical difference can be due to experimental variations in the PA set-up. An increase in PA signal magnitude was observed with increasing equivalent free dye concentration (5, 12 and 25 μM, Figure 5B) of the J-aggregates, which establishes the correlation between the total free dye concentration and PA signal intensity. The individual absorbance curves of all samples (JAAZ, N-JAAZ and RGD-JAAZ) at the above-mentioned concentrations are provided in Figure S23. The normalized absorbance of JAAZ and N-JAAZ shows a peak at 895 nm (Figure 5C, dashed lines) and the normalized PA profile matches the absorbance profile, with the PA signal peak at 890 nm (Figure 5C, solid lines). The slight shift of the PA signal maxima from the absorbance maximum can be attributed to slight offset between the wavelength setting and output wavelength of the tunable pulsed laser. The photoacoustic spectrum of RGD-JAAZ has a PA signal maximum at 890 nm, which is in the NIR-I region (Figure 5C), and optimally suited for deep-tissue imaging.

**Figure 5.**
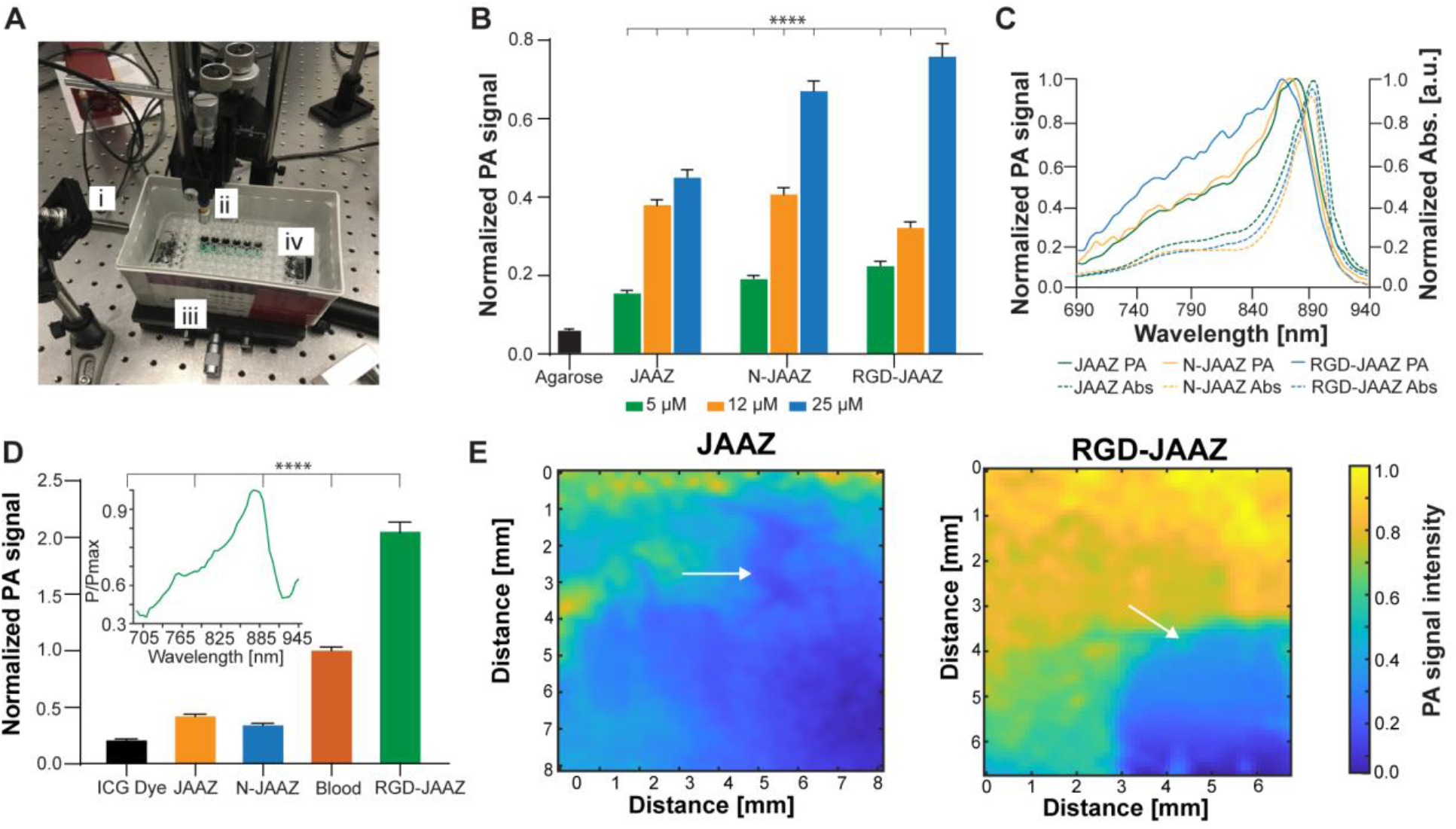
Photoacoustic properties of various JAAZ particles. (A) *In vitro* PAI set-up including laser source (i) transducer (ii) and reservoir containing deionized water (iii) to submerge the 96-well sample plate (iv) (B) PA signal of the different JAAZ particles at different concentrations. The normalization is performed with respect to 1:100 (v/v) dilution of India Ink. (C) Normalized absorbance (dashed line) and PA spectra (continuous line) for different JAAZ particles. (D) PA signal intensity of free ICG dye, and various JAAZ particles measured in sheep’s blood. Samples concentration was set at 20 μM and all values were normalized with respect to whole blood PA signal Individual data points are not shown for simplicity. Inset shows the PA spectra of RGD-JAAZ particles in whole blood. (E) 2D photoacoustic image of HeLa cells stained with 10 μM JAAZ and RGD-JAAZ particles showing photoacoustic signal normalized to maximum value. An area of the glass slide was scraped with a razor blade to remove cells before staining and the scraped region is indicated by an arrow in both figures. The entire scraped regions are highlighted in Figure S24A-B. Representative scale of PA signal amplitude is also given in the figure. (n=100 for B and D with data presented as mean±SD, p-values are calculated using one-way ANOVA; ****=p<0.0001; all P values are provided in Table S6).

To explore the possibility of using JAAZ particles as a contrast agent i*n vivo*, their photoacoustic signal amplitude was compared to that of whole blood. Short segments of 24-gauge PTFE tubing (containing approximately 100 μL of sample) were filled with 20 μM of ICG dye, JAAZ, N-JAAZ, RGD-JAAZ particles mixed in whole sheep’s blood. PA signals were recorded from each sample and the signal magnitude was normalized to the whole-blood PA signal magnitude (Figure 5D). The PA amplitude of 20 μM RGD-JAAZ was ~2.1x that of blood excited at 895nm, while 20 μM of free ICG dye was nearly ~0.03x the intensity of whole blood at this wavelength. The PA signal did not change in over 200 seconds when subjected to 2000 laser pulses (10 pulses/s) at 895 nm, which indicates that RGD-JAAZ particles are photostable (Figure S25A). Short segments of PTFE tubing were filled with 10 μM RGD-JAAZ and arranged in an ‘M’ pattern. The ‘M’ pattern was scanned to create a 2D PA image (Figure S25B). As a demonstration of cell labeling using JAAZ, 2D cultures of HeLa cells were stained with 10 μM JAAZ and RGD-JAAZ particles on a custom glass slide with a red silicone insert. Before staining with JAAZ and RGD-JAAZ, a specific area of cells was scraped away using a razor blade as shown in Figure S26A-B and the PA signal magnitude of the scraped region was used as a background to determine the level of non-specific binding of both particles. After washing the stained cells, a 35-MHz transducer was raster-scanned to produce a 2D PA image of the stained cells. The PA-color map indicates that the region where the cells were scraped away produced a weak PA signal that was less than 10% of the peak PA signal for cells stained with the particles (Figure 5E). When comparing regions with similar density of seeded (100,000 cells/mL) cells on the slide, RGD-JAAZ labeling resulted in a more uniform labeling of the cells as well as significantly stronger PA signals when compared to cells labeled with JAAZ (Figure 5E). Whereas the PA signal magnitude of the JAAZ stained cells decreased as the raster scan was completed (~3 hours), for the RGD-JAAZ stained cell culture, there was minimal to no diminution of the PA signal magnitude as the raster scan was completed (~3 hours). The direct maximum PA signal from each stained cell culture was compared for different laser intensities for RGD-JAAZ vs JAAZ with the signal from JAAZ stained cells being only half that of cells stained with RGD-JAAZ (Figure S26C). Our results indicate that the RGD-JAAZ particles can efficiently target and label integrin expressing HeLa cells taking advantage of the well-known binding affinity between RGD and the integrin receptors [42], as opposed to JAAZ particles without RGD, which do not bind to or label HeLa cells efficiently.

### 2.6. Pilot *In-vivo* imaging of two mice using RGD functionalized J-aggregates

To further demonstrate the clinical translation potential of the JAAZ particles, an *in vivo* pilot study was conducted on the commercially available small animal PAI system TriTom™ (Figure 6A)[43–45]. We first acquired multiwavelength images of microcuvette tubes containing 0.4 mM of ICG dye or RGD-JAAZ particles to evaluate the PA spectra of both contrast agents. The peak PA signal in the RGD-JAAZ tube was measured at 895 nm while the ICG dye tube had peaks at 740 nm and 780 nm (Figure S27). For *in vivo* imaging experiments, multispectral PAI were acquired in two female nu/nu nude mice at various timepoints (0, 10, 90 min) following tail-vein administration of ICG dye and RGD-JAAZ particles (Figure 6). A pre-injection baseline PA signal at 895 nm was also acquired for reference (Figure 6B). We found that while RGD-JAAZ and ICG dye demonstrate a similar biodistribution *in-vivo*, intravenous administration of RGD-JAAZ particles improved visualization of blood-rich tissues such as the liver and spleen for up to 90-minutes post-injection (Figure 6C and Figure S28A for PA images of the second mouse) compared to ICG. The improved visualization with RGD-JAAZ particles continued for up to 120-minutes post-injection as seen in Figure S28B. The achievable imaging depth was also improved by RGD-JAAZ particles demonstrated by the increased contrast-to-noise ratio (CNR) in blood vessels as deep as 5 mm from the surface of the skin compared to ICG dye 90-minutes post-injection (Figure 6D and Table 1). The circulation half-life of RGD-JAAZ particles was 10 times greater than ICG dye (Table 1). Additionally, the molecular maps of oxyhemoglobin, deoxyhemoglobin, and RGD-JAAZ particles show the biodistribution of the contrast agent is restricted to the vasculature and blood-rich tissues such as the liver and spleen (Figure 6E). Molecular maps were generated from a linear spectral unmixing of the multiwavelength PA images using previously reported spectra for hemoglobin and the measured spectra of the RGD-JAAZ particles at their injection concentration [45,46].

**Figure 6.**
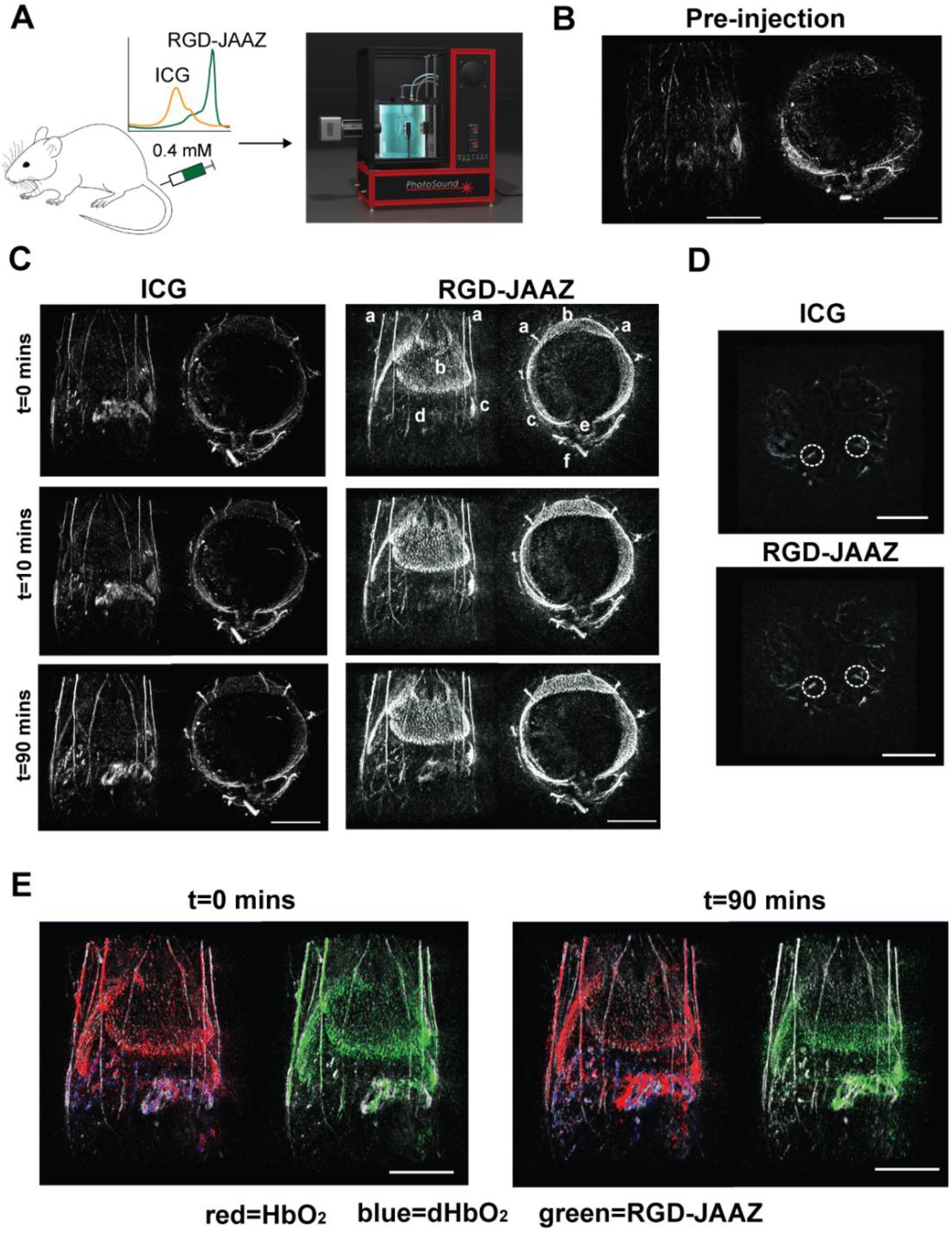
*In vivo* imaging using RGD-JAAZ particles. (A) Schematic and photograph of the 3D PAI acquisition on the TriTom™ imaging system. (B) Coronal and axial slabs of the baseline PA signal acquired at 895 nm. (C) Representative coronal and axial slabs of the ICG (left) and RGD-JAAZ particle (right) injections acquired at the peak excitation wavelength (i.e., 800 and 895 nm, respectively) over time. The displayed dynamic range for all images was set based on the baseline scan acquired at the same wavelength. (Legend): a. superficial thoracic vessels, b. liver c. spleen d. intestines e. thoracic vertebrae f. CuSO_4_ fiducial tube. (D) Axial slabs encompassing the iliac arteries (white ROI) used to calculate the achievable imaging depth for ICG and RGD-JAAZ particles 90-minutes post-injection. (E) Coronal slabs of oxyhemoglobin (red), deoxyhemoglobin (blue), and RGD-JAAZ particles (green) spectral unmixing overlaid on the 800 nm excitation scan of anatomy (white) at 0- and 90-minutes post-injection. Scale bars for all images are 10 mm. (n= 2 female nu/nu nude mice)

**Table 1:** Contrast-to-noise ratio and circulation half-life of ICG and RGD-JAAZ particles.

| Injection | Contrast-to-noise ratio | Circulation half-life (mins) |
| --- | --- | --- |
| ICG | 1.59 | 1.19±0.45 |
| RGD-JAAZ | 2.42 | 10.26±1.24 |

n=2. Data is represented as mean±SD.

## 3. Conclusion

In this study, we designed and validated a simple and robust protocol for the synthesis of functionalized NIR-PAI contrast agent with tunable sizes ranging from 1.2 μm to 230 nm, which allows photoacoustic imaging at an excitation wavelength of 895 nm. As an added advantage over previously reported ICG J-aggregates [29], the modified J-aggregates synthesized in our study do not require size selection or further downstream processing. The particles are also stable in physiological conditions and can be easily dried and resuspended for long-term storage.

We successfully demonstrated direct functionalization of the azide groups using efficient and reliable copper-free click chemistry, which allow for further conjugation with biomolecules and targeting moieties such as streptavidin and RGD onto the surface of JAAZ and N-JAAZ. Our functionalized particles exhibited a strong PA signal even at concentrations as low as 5 μM equivalent free dye and showed nearly 2.1x greater PA magnitude in comparison with whole blood. RGD-JAAZ demonstrated the cell targeting capabilities of our platform with a significantly higher PA magnitude compared to JAAZ stained HeLa cells. The *in vivo* 3D PAI images showed that RGD-JAAZ particles can be visualized with a CNR of 2.42 in blood vessels as deep as 5 mm from surface of the skin and generate strong and quantifiable PA signals in blood-rich tissues such as the liver and spleen for up to 90-minutes post-injection in comparison with ICG. At 90 minutes of injection, in the RGD-JAAZ unmixed images, high intensity signals of these particles are clearly visible in liver and spleen over hemoglobin and deoxyhemoglobin. Hence, these results demonstrate the potential use of RGD-JAAZ as a promising contrast agent platform for future applications such as tumor margin staining, [29] imaging diseased tissue [28] and lymph node biopsy. [47]

## 4.1. Experimental Section

### 4.1.1. Materials

Indocyanine Green (ICG) and ICG-Azide were purchased from AAT Bioquest. Dibenzylcyclooctyne-PEG4-Biotin (DBCO-PEG4-Biotin) was acquired from Jena Biosciences. The streptavidin and the Cyclo-Arg-Gly-Asp-D-Lys(Biotin) (RGD-Biotin) were purchased from Peptides International. The Amicon Ultra-0.5 centrifugal filters, potassium chloride (KCl), agarose, molecular biology grade water, penicillin, and Bicinchoninic Acid (BCA) Kit were acquired from Sigma-Aldrich. Triton X-100, Tween-20, Dimethylsulfoxide (DMSO) and glycerol were purchased from Fischer Scientific. Phosphate-Buffered Saline (PBS, 10X solution at pH 7.4) was acquired from ThermoScientific. A Low melt Agarose was purchased from IBI Scientific. Molecular biology grade water was purchased from Quality Biological. The low-volume quartz glass cuvettes and the folded capillary zeta cells were purchased from Malvern Panalytical. 1X Dulbecco’s Modified Eagle’s Medium was purchased from Corning Inc. and Fetal Bovine Serum (FBS) were purchased from VWR Life Sciences. India Ink and whole sheep blood containing anticoagulant were purchased from Hardy Diagnostics. Eagle’s Minimum Essential Media (EMEM) was purchased from ATCC. Cell Proliferation Kit I (MTT) was purchased from Millipore Sigma with active ingredient 3-(4,5-dimethylthiazol-2-yl-2,5-diphenyltetrazoliumbromide (MTT).

### 4.2. Methods

#### 4.2.1. Synthesis of ICG-JA, JAAZ and N-JAAZ particles

ICG and ICG-azide stock solutions were respectively prepared at a concentration of 5 mg/ml (6 mM) in water and 10 mg/ml (12 mM) in DMSO and stored at −20°C. To assemble the ICG-JA, ICG dye was prepared at 1 mM in water complemented with 20 mM of KCl and incubated at 60°C for 20 hours. For JAAZ particles, ICG and ICG stock solutions were mixed to reach a final ICG concentration of 1 mM and different molarity of ICG-azide in water with 20 mM KCl. The mixes were incubated for 20 hours at 60 °C. For JAAZ particles in the 600-700 nm range, the reaction solution of 1:10 ICG-azide:ICG molar ratio was incubated in 1 mM KCl for 20 hours at the same temperature as mentioned above to a final concentration of 1 mM total dye. To make N-JAAZ particles of size ~200 nm, 250 μM total dye solution was incubated in 0.1 mM KCl for 14 hours at 60 °C. A complete summary of the total dye concentration at 1:10 ICG-azide:ICG molar ratio, KCl molarity and time required for the formation of JAAZ particles is given in Figure S18. Before further use, ICG-JA, JAAZ, and N-JAAZ samples were filtered using 0.5 ml Amicon centrifugal filters (100 kDa cutoff) to remove the non-aggregated free monomers of ICG and exchange the buffer. Filtration was carried out by centrifugation at 4000*g* for 3 minutes at room temperature. Three cycles of filtration in water were used to ensure complete removal of unreacted dye. The percentage of samples recovered after filtration was calculated by:

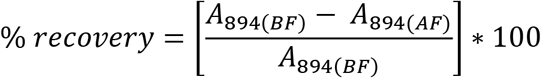

Where, *A*_894(*BF*)_ is the absorbance intensity at 894 nm before filtration (BF) and, *A*_894(*AF*)_ is the absorbance intensity after filtration.

#### 4.2.2. Characterization of JAAZ particles

Samples were scanned from 440 nm to 990 nm with 2 nm step size using a PowerWavex microplate spectrophotometer by Bio-Tek Instruments Inc. The amount of free dye in all J-aggregate samples was determined by incubating the formed J-aggregate particles in 1% Triton X-100 for 10 minutes at 37°C and scanning the samples from 400-990 nm. A standard curve of a 1:10 molar solution of ICG-azide:ICG dye was created to determine the free dye concentration in the dissociated J-aggregate samples. Fluorescence intensity of different concentrations of free ICG dye and J-aggregates was recorded using a Tecan Safire^2^ fluorescence microplate reader. Both free ICG dye and J-aggregate particles were excited at 720 nm with an emission at 770 nm. The fluorescence emission spectrum was recorded from 770 to 850 nm with a step size of 2 nm. The hydrodynamic diameter of the formed ICG-JA and JAAZ particles was measured using a Malvern Zetasizer Nano ZS instrument. The size measurements were taken using low-volume quartz glass cuvettes at 25 μM of equivalent free dye concentration. The zeta potential (surface charge) of all samples was determined using the Malvern Zetasizer Nano ZS using DTS-1070 folded capillary zeta cells. The measurements for zeta potential were taken at an equivalent free dye concentration of 15 μM and a volume of 700 μl in 0.1X PBS at pH 7.4. A 2% (w/v) gel was prepared using high strength agarose in 1X TAE buffer for electrophoresis. A square window of 6 x 6 cm was cut in the 2% gel. A 0.1 % (w/v) low melting agarose gel prepared in 1X TAE buffer was poured in the square window and let to polymerize at 4°C overnight. 30 μl of 500 μM JAAZ and N-JAAZ samples were mixed with an equal volume of glycerol and loaded into the wells. The gel was run at 120 V for approximately 10 minutes. For scanning electron microscopy, 2 μl of the JAAZ or the N-JAAZ samples were deposited on a clean mica surface and dried overnight under vacuum. SEM images were collected using a JEOL JSM-IT500HR InTouchScope™ at an accelerating voltage of 15 kV and a working distance of 10 mm. EDX spectra were collected using Octane Elect EDS System at an accelerating voltage of 15 kV and a resolution of 127 eV. ImageJ [48] was used to determine the size distribution of the particles in the obtained SEM images.

#### 4.2.3. Functionalization of JAAZ particles

500 μM of JAAZ or N-JAAZ particles with 1:10 ICG-azide:ICG molar ratio were labelled with DBCO-PEG_4_-Biotin at a concentration 10 times greater than the estimated number of azide groups available (Table S2). The samples were incubated and mixed in an incubator shaker overnight at room temperature protected from light. To remove excess DBCO-Biotin, the samples were filtered as mentioned above. These biotin functionalized samples (Bio-JAAZ) were diluted to an equivalent of 100 μM of free dye and incubated with streptavidin at 5X concentration of the available biotin sites (Strep-JAAZ). The samples were kept at 4°C for two hours to ensure attachment of streptavidin to the available biotin sites. Excess streptavidin was removed by filtration. The Bio-JAAZ and the Strep-JAAZ samples were characterized using the procedures mentioned in the characterization section. The Strep-JAAZ samples were labelled with RGD-Biotin at 10X molar concentration and incubated for two hours at 4°C. Excess RGD-Biotin was removed by filtration using the same procedure as mentioned before. The labelled samples were named RGD-JAAZ. The RGD-JAAZ samples were characterized using the procedures mentioned in the characterization section. Purified Strep-JAAZ samples were diluted to an equivalent of 5 μM of equivalent free dye concentration. BCA working reagent was made following the instructions provide by the supplier. 25 μl of 5 μM samples are incubated in 200 μl of BCA working solution and the amount of streptavidin attached is calculated using a BSA standard curve.

#### 4.2.3. Hemotoxicity, Cytotoxicity and stability in whole blood of JAAZ particles

Whole sheep’s blood was centrifuged at 1000*g* for 2 minutes at 4 °C for 2 minutes to remove any coagulated blood. 80 μl of the centrifuged blood was mixed with 20 μl of samples by gently pipetting up and down and incubated at 37 °C for 30 minutes. 10 μl of treated blood was pipetted onto a clean glass slide and another glass slide was scraped across its entire length to create a blood smear. Microscopy images were acquired using Zeiss AxioVert 200 microscope. Hela cells were used for the cytotoxicity experiment of RGD-JAAZ, N-JAAZ, and JAAZ. The cells were seeded at 5 x10^4^ cells/well in a 96-well plate incubated in augmented EMEM at 37°C and 5% CO2 for 24 hours. Cells were then cultured with 10μM of RGD-JAAZ, N-JAAZ, or JAAZ. 10μL (final concentration of 0.5 mg/mL) of the standard 3-(4,5-dimethylthiazol-2-yl-2,5-diphenyltetrazoliumbromide (MTT) was added to each well of the 96-well plate and incubated for 4 hours. Then 100 μL of solubilization solution was added to each well and the plate was incubated overnight. The absorbance of each well was tested by a microplate (ELISA) reader (accuSkan FC FisherBrand, Waltham, MA, USA) measuring the absorbance of the formazan product between 550 and 600 nm. A positive control group was tested with an equivalent mixture of saline mixed with culture medium.

80 μl of whole blood was incubated with JAAZ, N-JAAZ and RGD-JAAZ samples respectively, at 37 °C. The final concentration of all samples in whole blood was 10 μM. At 0, 2, 4, 6, 8, 16 and 24 hour, 60 μl of the incubated sample for each condition was scanned from 440-990 nm with 2 nm steps using a microplate spectrophotometer.

#### 4.4.4. Characterization of the photoacoustic properties of JAAZ particles *in-vitro*

The 96-well plate containing samples embedded in agarose was secured to the bottom of a plastic water reservoir. The reservoir was filled with deionized water, completely submerging the samples. For comparison of smaller volumes, 2-inch segments of 24-gauge PTFE tubing (~100 μL) were filled with various samples. The tubes were placed in a clear acrylic chamber submerging the samples in deionized water. A single element piezoelectric 35MHz (PI35, Olympus, Massachusetts, USA) or 5MHz (V326, Olympus, Massachusetts, USA) focused transducer was fixed at 12mm or 45mm above each sample with their functional element submerged in deionized water. Optimization of positioning and photoacoustic signal from each sample was done though movement of a fine x-y axis stage, moving above each sample with a fixed transducer and pulsed laser. Excitation of the sample was done obliquely by a wavelength-tunable (690-950 nm) pulsed laser designed for photoacoustic imaging (Phocus Mobile, Opotek) at the peak absorption wavelength for JAAZ particles (895nm) for magnitude comparison. The photoacoustic signal recorded from the transducer is sent to a 20/40db amplifier (HVA-200M, Femto) from there the signal is recorded through a lock-in amplifier (Zurich Instruments, Zurich, Switzerland) triggered on each laser pulse using a photodiode placed near the laser source. For each measurement, a 10 second recording (100 waveforms) was taken for each sample (or at each wavelength, for PA spectrum generation). The absolute amplitude of each waveform was calculated, averaged, and normalized to the signal recorded from India ink or whole sheep’s blood containing anticoagulant. For cell targeting and photoacoustic imaging, Hela cells were cultured in EMEM augmented with 10% Fetal Bovine Serum, and 1% Penicillin and kept in an incubator at 37°C and 5% CO2. HeLa cells were seeded using 0.25% (w/v) Trypsin-0.53 mM EDTA solution to glass slides at a concentration of 1.0×10^5^ cells/mL. The Hela cells were stained with 10 μM JAAZ or RGD-JAAZ for 20 minutes in an incubator at 37°C and 5% CO2. Short segments of 24-gauge PTFE tubing (~100 μL) were filled with 10 μM RGD-JAAZ and arranged on a glass slide. The slides were then placed under a single element piezoelectric 35 MHz (or 15 MHz) focused transducer fixed at 12 mm (or 2 inches) above the surface of the slide. A wavelength-tunable (690-950 nm) pulsed laser designed for photoacoustic imaging (Phocus Mobile, Opotek) was obliquely shown over the slide. The peak absorption wavelength for the JAAZ (895 nm) was used to excite the sample and the photoacoustic signal recorded from the transducer was sent to a 20/40 dB amplifier (HVA-200M, Femto) from there the signal was recorded through a lock-in amplifier (Zurich Instruments, Zurich, Switzerland) triggered on each laser pulse using a photodiode placed near the laser source. The sample was then raster scanned through movement of the stage at 100 μm steps (with fixed laser source and transducer) mounted on motorized translation stages (Thorlabs). Absolute signal magnitude was then plotted in 2D space to create the image. A simple Matlab image gaussian filter was then used to smooth the image.

#### 4.2.5. *In-vivo* photoacoustic imaging using RGD-JAAZ particles

The phantom and *in vivo* photoacoustic imaging studies were performed using the TriTom™ imaging platform (PhotoSound Technologies, Inc.) and 20 Hz nanosecond-pulsed OPO laser (PhotoSonus, EKSPLA) located at MD Anderson Cancer Research Center in Houston, TX. The TriTom is a commercially available multimodal imaging technology capable of high-resolution, 3D multispectral photoacoustic imaging in the whole body of small animal models[43]. To prepare for imaging, the TriTom™ imaging chamber was filled with degassed, deionized water at room temperature for phantom experiments or heated to 36.0 ± 0.5°C for animal studies. In both studies, free ICG dye was diluted to 0.4 mM from a stock solution and the same concentration of RGD-JAAZ particles was prepared using the methods mentioned in the above sections. The size of the RGD-JAAZ particles was 1.24±0.24 μm.

The sample phantom was constructed using 810 μm inner diameter PTFE microcuvette tubes filled with a 50 μL volume of water, PBS, free ICG dye, or RGD-JAAZ particles. The four sample tubes were secured in a phantom holder that was then mounted in the TriTom and lowered into the imaging chamber. A series of single-wavelength 3D PA scans were acquired using excitation wavelengths from 700 to 900 nm in 10 nm steps with an additional scan at 895 nm. During each scan, the sample phantom was rotated 360° while acquiring 760 ± 5 frames of PA data. The acquired PA data were reconstructed into 30×30×30 mm volumes with a voxel size of 0.1 mm using a filtered back-projection method [49]. All visualization and analysis of the reconstructed volumes was performed in 3D Slicer. To determine the PA spectra of the ICG and RGD-JAAZ samples measured with the TriTom™, a 5 mm vertical section in the center of each microcuvette tube was manually segmented and the maximum PA signal intensity was calculated for all wavelengths.

A pilot study consisting of two female nu/nu nude mice (Charles River Laboratories) was conducted on the TriTom to compare the RGD-JAAZ particles of size 1.24±0.24 μm to ICG dye *in-vivo*. During each imaging session, the animal was anesthetized with isoflurane and a saline-filled catheter was inserted in the tail vein and secured. The animal was then transferred to a mouse holder and mounted in the TriTom imaging chamber so that the liver, spleen, and abdominal region of the mouse were visible. First, a baseline series of 3D PAI data was acquired with 700, 750, 800, 850, 875, 885, and 895 nm excitation wavelengths. A 100 μL bolus of 0.4 mM free ICG dye was then injected via the tail vein catheter followed by a 50 μL saline flush and the same set of PAI data was collected at 0, 10, 30, 60, and 90-minutes post-injection. After the final imaging time point, the PA scan acquired at 800 nm was reconstructed and compared to the pre-injection scan to confirm that the ICG dye had completely cleared circulation. A matching 100 μL bolus of 0.4 mM RGD-JAAZ particles and 50 μL saline flush was then administered and the 3D PAI imaging sequence was repeated with an additional time point at 120 minutes post-injection. Each 3D PAI scan was then reconstructed into a 30×30×30 mm volume with a voxel size of 0.1 mm using a filtered back-projection method. The displayed dynamic range was determined using the baseline scans and set for all 3D reconstructions and axial slices.

To estimate the circulation half-life of RGD-JAAZ particles compared to monomeric ICG, four superficial blood vessels in the thorax region were manually segmented for the 0, 10, 30, and 60-minute post-injection volumes. Half-life estimation was performed using the 800 nm and 895 nm excitation scans (i.e., peak absorbance wavelength) for ICG and RGD-JAAZ particles, respectively. The baseline PA signal in each vessel was subtracted and then averaged for each time point. An exponential function fit to the data in each animal was then used to calculate the circulation half-life of ICG or RGD-JAAZ [24].

The achievable imaging depth of RGD-JAAZ particles and monomeric ICG was assessed by comparing the CNR in the iliac arteries from the baseline and 90-minute post-injection scans. Three axial slices containing both blood vessels at varying depths from the skin were identified using the 3D reconstructed volumes. The vessels and background signal (i.e., the area surrounding the animal) were manually segmented and the shortest distance from the center of each vessel to the skin was measured. CNR was then calculated for each artery as:

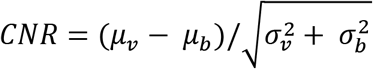

Where μ and σ are the averages and standard deviation of the vessel and background, respectively. The blood vessel was considered detectable at the measured imaging depth if the calculated CNR was ≥ 2. Finally, multispectral PAI (700, 750, 800, 850 and 895 nm) was performed to spectrally unmix the endogenous PA signal from oxyhemoglobin, deoxyhemoglobin from the RGD-JAAZ signal. The unmixed volumes were then overlaid on the reconstructed 800 nm scan for anatomical reference.

##### Statistical Analysis

All bar graphs are expressed as the mean ± standard deviation. Statistical analyses were performed using the unpaired, two-tailed Student’s t-test and One-Way Analysis of Variance (ANOVA) with a Tukey post-hoc test using the software GraphPad Prism 9.0.2. Data was deemed statistically significant if the *p*-value was <0.05. Absolute PA signal amplitude is defined as the sum of the absolute minimum and maximum peak PA signal.

## Supporting Information

The supplementary information is provided with the article.

## Data Availability

Data is available upon request.

## Ethical Statement

All animal studies were approved by the Institutional Animal Care and Use Committee at MD Anderson Cancer Center. IUCAC protocol number 00001779-RN01, “Evaluations of Tumor Models with Multispectral Optoacoustic Tomography.”

## Author Contributions

S.S. and G.G. contributed equally to the manuscript. S.S. synthesized all the J-aggregate samples, performed the characterization and the conjugation experiments, and analyzed the data. G.G., P.C., helped by S.S., and R.V. designed the photoacoustic experiments. G.G. performed the photoacoustic experiments and analyzed the data. C-H.H, J.B. and L.S.C performed additional characterization experiments and data analysis. D.J.L. performed the phantom and *in-vivo* experiments along with the associated data analysis. J.L.M. performed the SEM imaging and assisted with data analysis. R.V. designed and supervised the study and interpreted the results. S.S., G.G., D.J.L., and R.V. wrote the manuscript. All authors commented on and edited the manuscript.

## Conflict of Interest

S.S, P.V.C. and R.V. hold a provisional patent covering the work presented in the manuscript. All other authors declare no competing interest.

## Acknowledgement

S.S. gratefully acknowledges George Mason University, Office of Provost and Executive Vice President for the Summer 2019 Provost Research fellowship. L.S.C thanks for his support through the summer Undergraduate Research Scholars Program (URSP) from the Office of Student Scholarship, Creative Activities and Research at George Mason University. We thank Dr. Mikell Paige, Associate Professor in the Department of Chemistry and Biochemistry at George Mason University to allow use of instruments in his lab. We acknowledge Sergey Ermilov, Chief Executive Officer at PhotoSound Technologies Inc. for his helpful insights in the *in vivo* studies. We also thank Dr. Mark D. Pagel, Professor, Department of Cancer Systems Imaging, University of Texas MD Anderson Cancer Center and his research team for providing the mice used in the *in vivo* studies.

