## Supplementary Material for "Size-tunable ICG-based contrast agent platform for targeted near-infrared photoacoustic imaging"

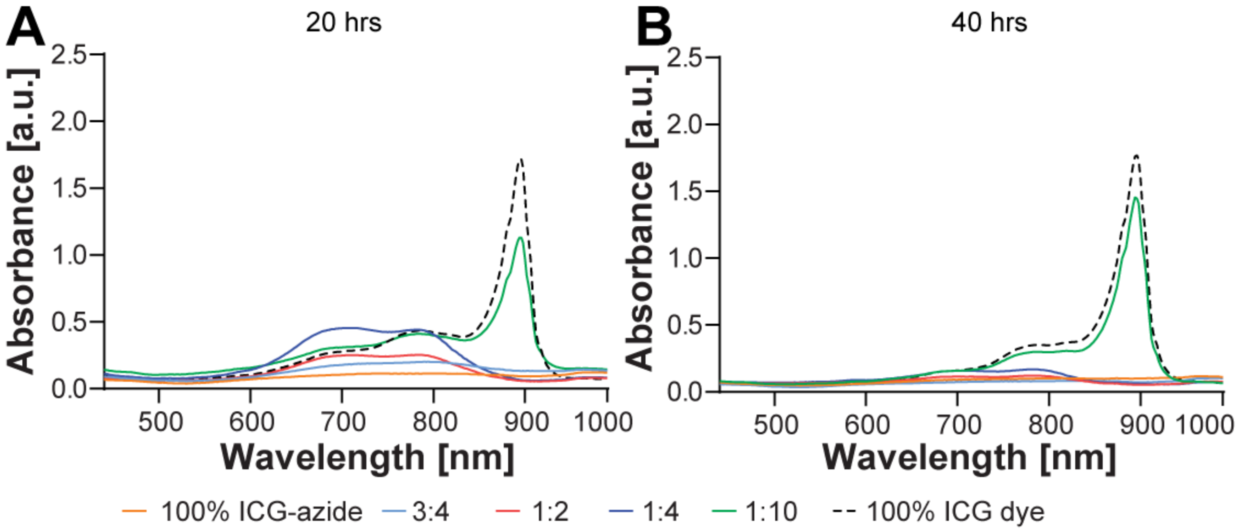


**Figure S1. Lack of formaation of JAAZ in water**. (A) Absorbance spectrum of 100% ICG-azide, 100% ICG dye and different molar ratio solutions of ICG-azide:ICG made in water after 20 hours of incubation (n=2). (B) Absorbance readings after 40 hours of incubation (n=2). Absorbance measurements taken at 25 $\mu$M equivalent free dye concentration.

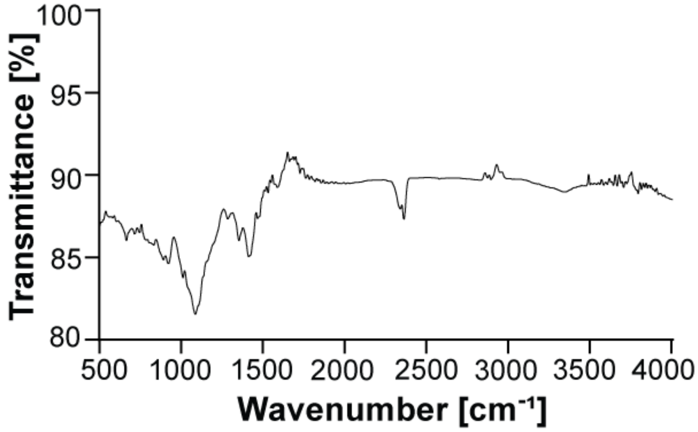


**Figure S2.** FT-IR spectrum of ICG-JA particles with 1:10 ICG-azide:ICG molar ratio formed in water (n=2) at 60 °C. The lack of a peak between 2141-2100 $cm^{-1}$ indicates that no azide was incorporated into the J-aggregates.

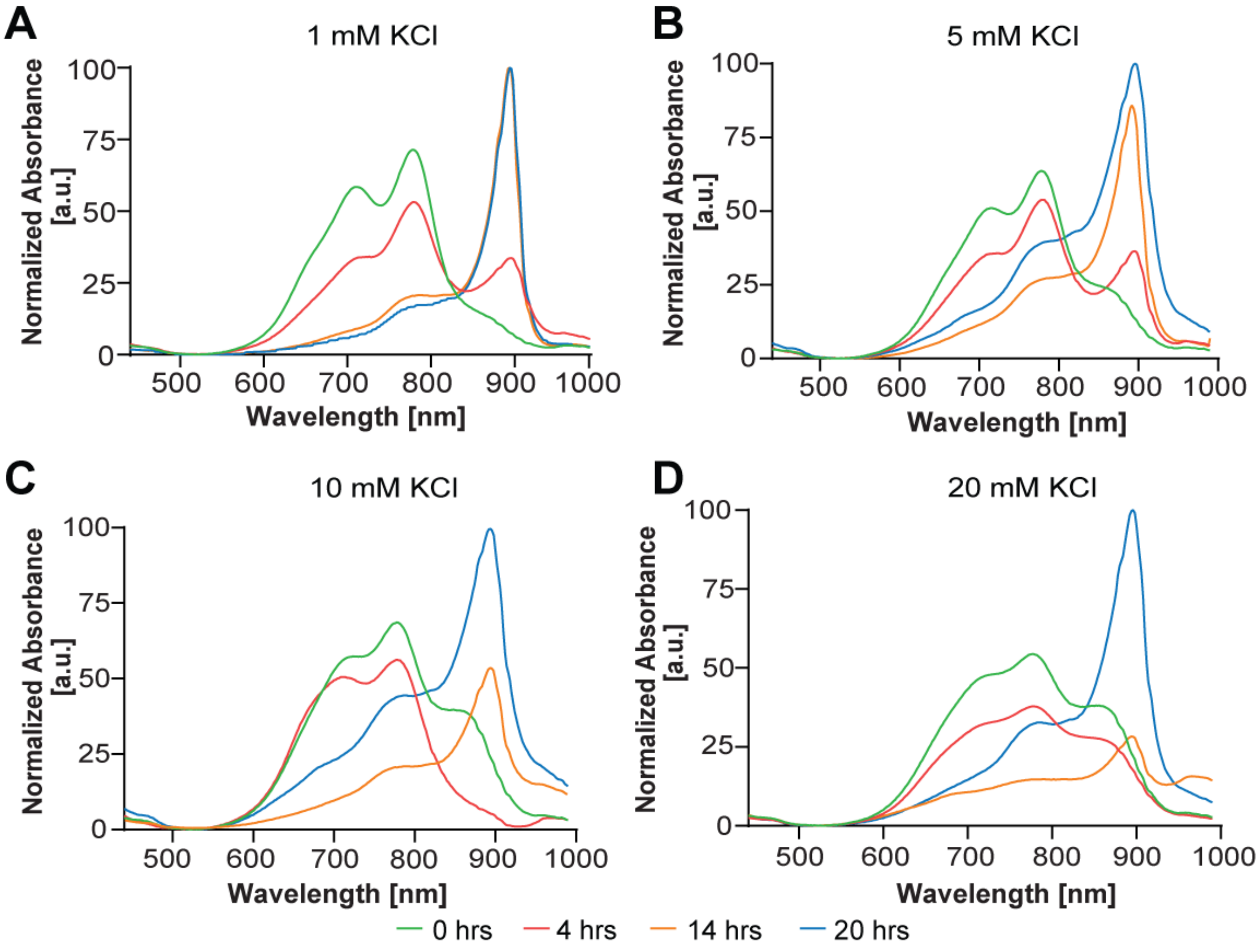


**Figure S3. Formation of JAAZ in KCl.** (A-D) Absorbance spectrums of 1:10 ICG-azide:ICG molar solution at dye concentration of 1 mM in different KCl concentrations of 1 mM (A), 5 mM (B), 10 mM (C) and 20 mM (D) incubated for 20 hours at 60 °C (n=3).

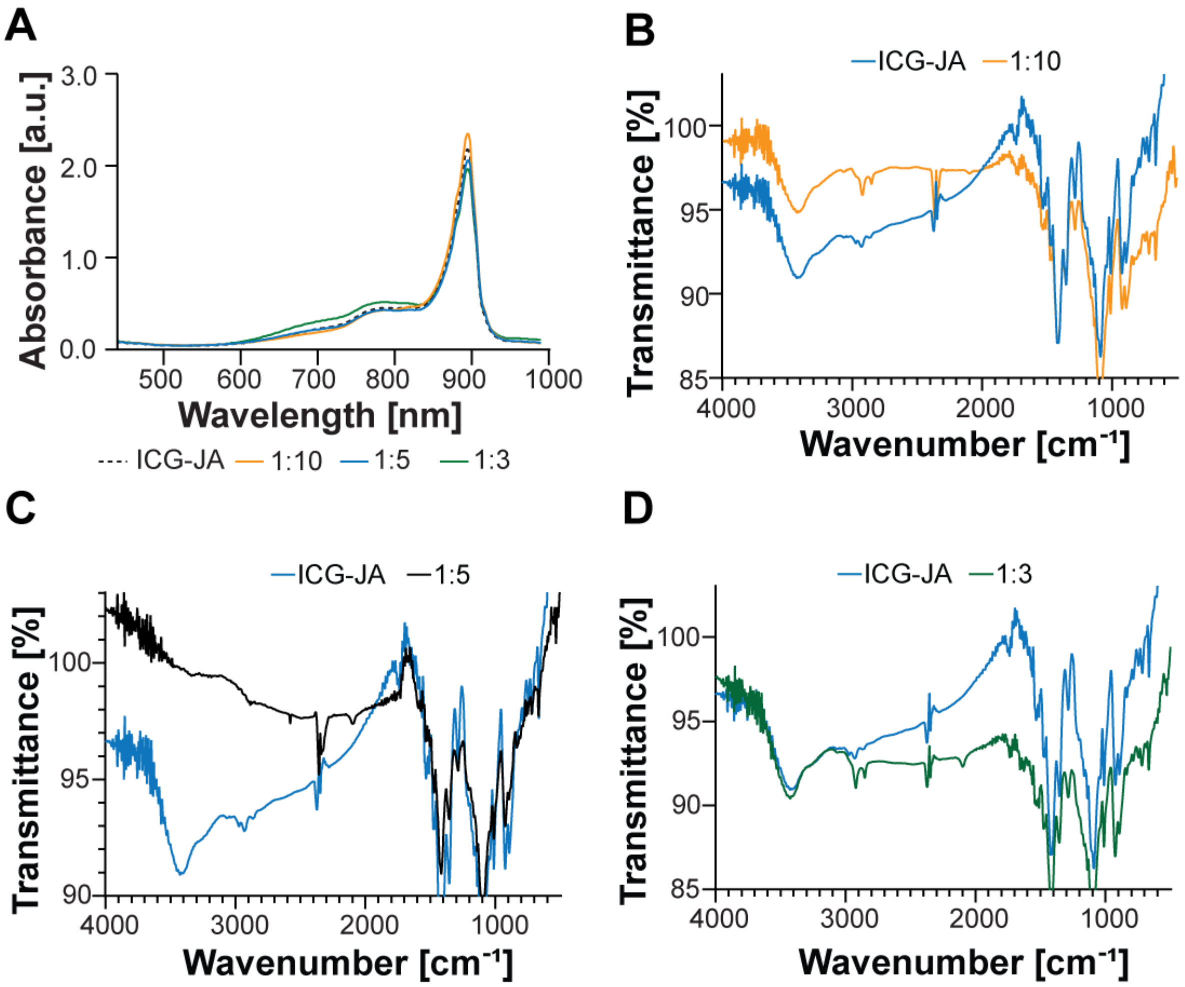


**Figure S4. UV-Vis-NIR and FT-IR spectrum of different molar ratio JAAZ particles**. (A) UV-Vis-NIR spectrum of JAAZ particles formed with different molar solutions of ICG-azide:ICG (1:10, 1:5, 1:3) to a final dye concentration of 1 mM. Absorbance readings taken at an equivalent free dye concentration of 50 $\mu$M (n=3). (B-D) FT-IR spectrum of JAAZ particles with (B) 1:10 (orange), (C) 1:5 (black), and (D) 1:3 (green) molar ratio of ICG-azide:ICG. The FT-IR spectrum of the control is ICG-JA particles is depicted in blue (n=1).

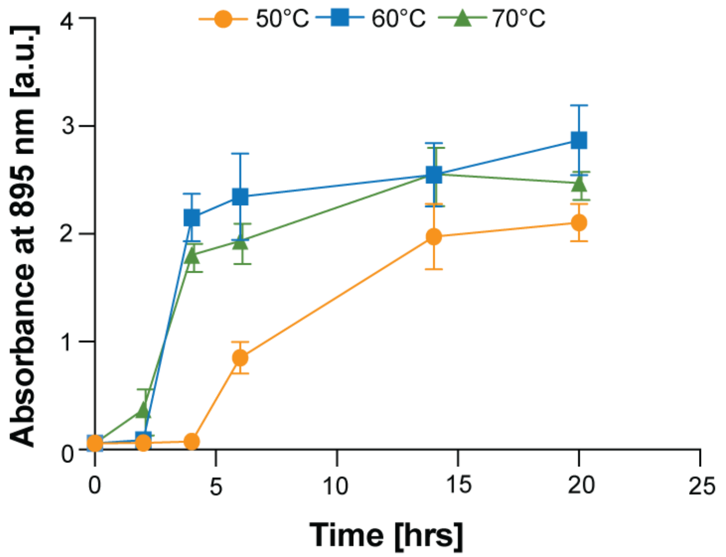


**Figure S5. Rate of formation of JAAZ particles at different temperatures** Absorbance intensity at 895 nm of the JAAZ particles formed at 50 °C (orange), 60 °C (blue) and 70 °C (green). The final dye concentration in all samples is 1 mM

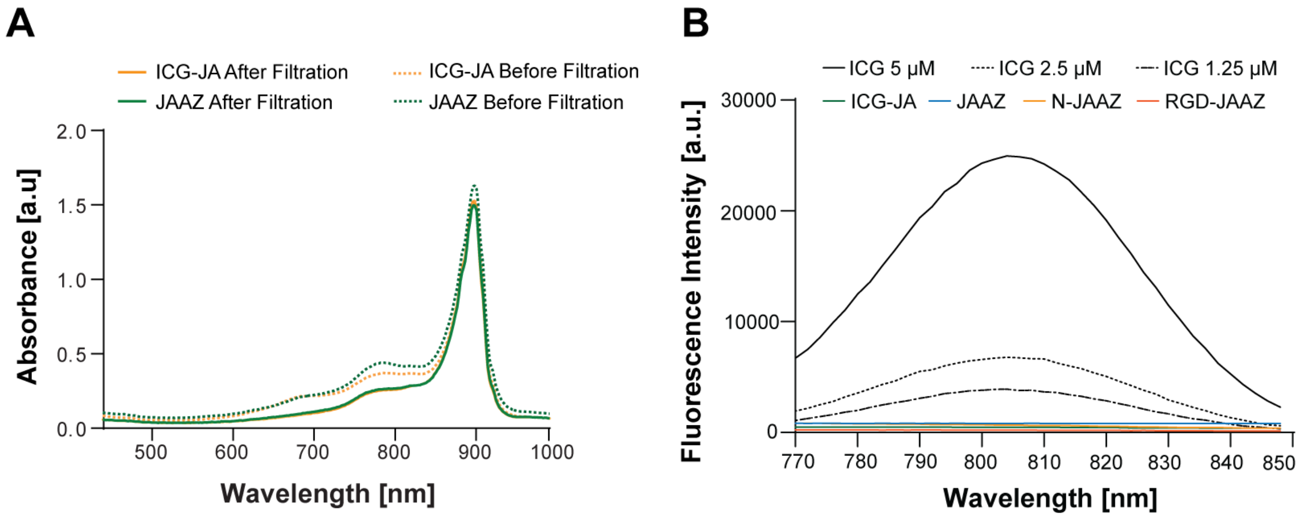


**Figure S6. Filtration and Fluorescence of ICG-JA and JAAZ**. (A) Absorbance spectra of the formed ICG-JA and JAAZ before and after filtration showing a clear peak at 895 nm, typical of ICG J-aggregates. Absorbance measurements taken at 25 $\mu$M equivalent free dye concentrationfor the filtered samples (n=2). (B) Fluorescence emission curves of free ICG dye at concentrations 5, 2.5 and 1.25 $\mu$M and those of ICG-JA and JAAZ particles. Fluorescence emission intensity is absent in both ICG-JA and JAAZ particles.

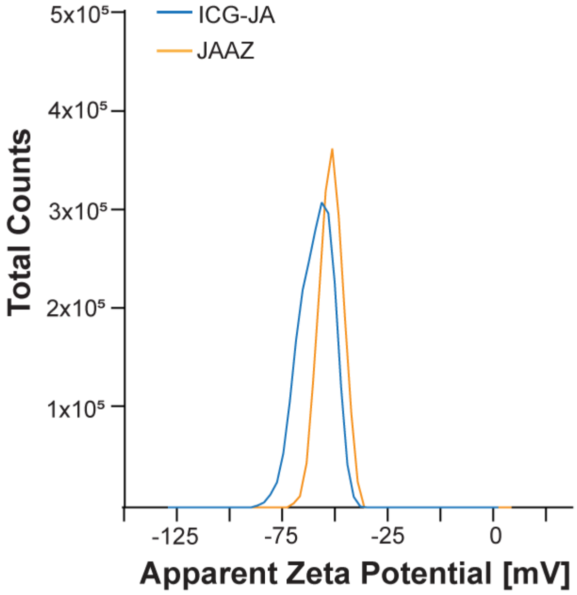


**Figure S7. Zeta potential distribution of ICG-JA and JAAZ**. Charge distribution profile of ICG-JA and JAAZ centered at -61 mV for ICG-JA and -52 mV for JAAZ (n=3).

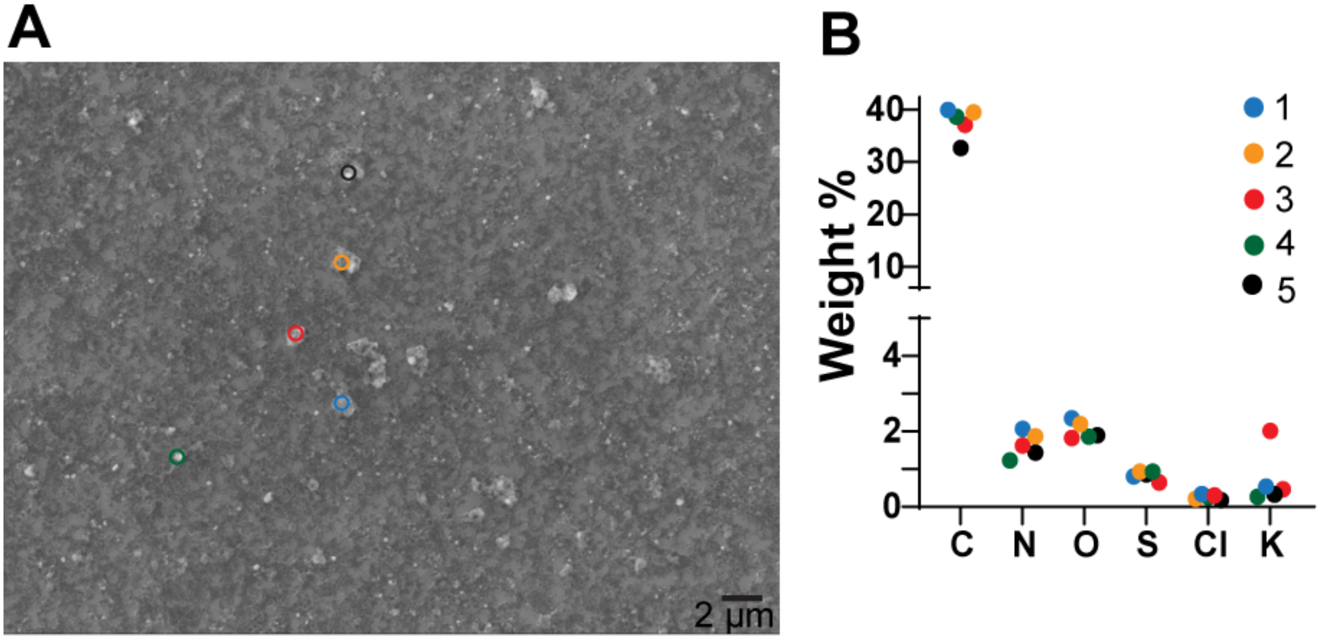


**Figure S8. SEM image and EDX analysis of JAAZ particles**. (A) SEM image of JAAZ particles prepared at 1 mM dye concentration and 20 mM KCl. Size measurements of these JAAZ particles using DLS showed an average of 900 nm. (B) EDX analysis showing weight percentage of carbon (C), nitrogen (N), oxygen (O), sulfur (S), potassium (K) and chloride (Cl) of five different JAAZ particles as marked in image a**.**

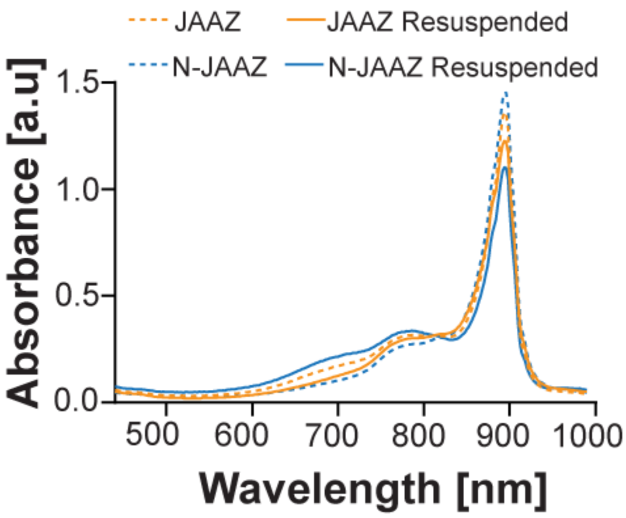


**Figure S9. Drying and resuspension of JAAZ particles**. Absorbance curves of JAAZ and N-JAAZ particles before drying and after resuspension (n=2). Absorbance measurements taken at 25 $\mu$M of equivalent free dye concentration.

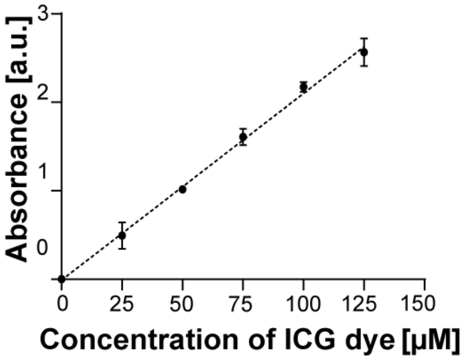


**Figure S10. Standard Curve of ICG dye.** Standard Curve of ICG dye monomer or free ICG dye used to determine the equivalent free dye concentration in disassociated ICG-JA and JAAZ particles.

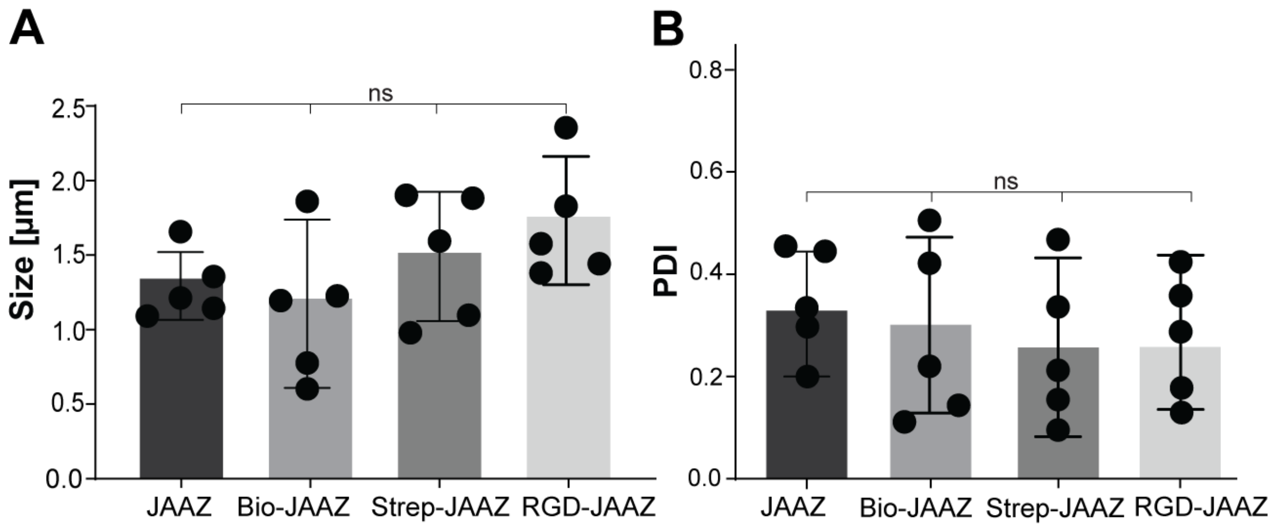


**Figure S11. Size and PDI of Streptavidin functionalized JAAZ**. (A) The size of the different biomolecule modified forms of JAAZ (data presented as mean±SD, n=5, p-values are calculated using one-way ANOVA, p=ns, not significant; all P values are provided in Table S7). (B) The PDI of JAAZ, Bio-JAAZ, Strep-JAAZ and RGD-JAAZ was 0.36, 0.33, 0.27 and 0.26 respectively. (data presented as mean±SD, n=5, p-values are calculated using one-way ANOVA, p=ns, not significant; all P values are provided in Table S7).

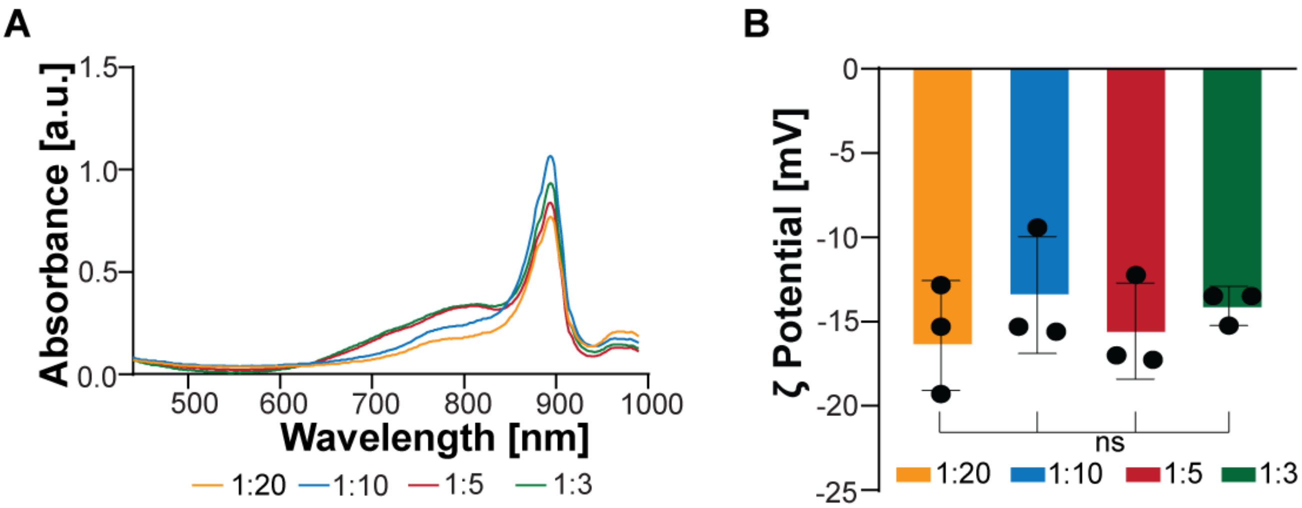


**Figure S12. UV-Vis-NIR spectrum and zeta potential of different molar ratio Strep-JAAZ particles.** (A) Absorbance spectrum of different ICG-azide:ICG molar ratio JAAZ particles modified with streptavidin (n=3). Absorbance measurement taken at 25 $\mu$M equivalent free dye concentration. (B) The zeta potential of 1:20, 1:10, 1:5 and 1:3 Strep-JAAZ samples are -16.4, -13.3, -15.5 and -14.1 respectively (data presented as mean±SD, n=3, p-values are calculated using one-way ANOVA, p=ns, not significant; all P values are provided in Table S8).

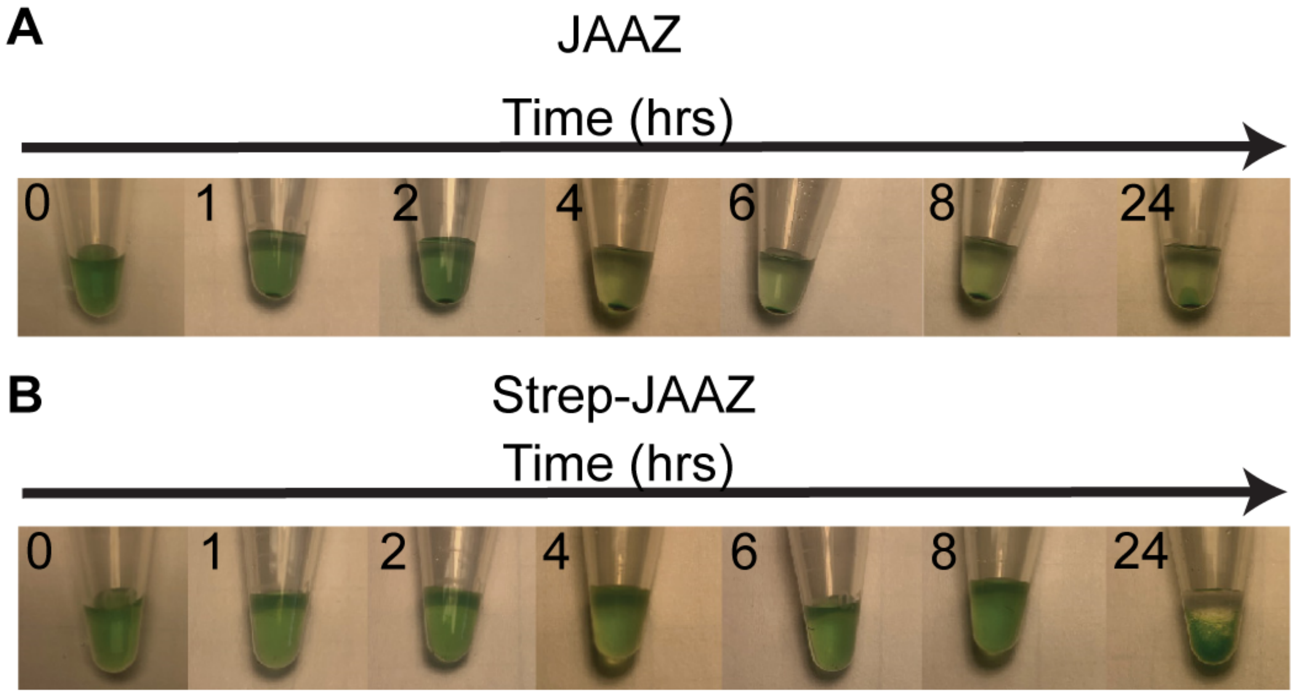


**Figure S13. Sedimentation of JAAZ and Strep-JAAZ for 24 hours.** (A) Pictures of JAAZ sample at 0, 1, 2, 4, 6, 8 and 24 hours of sedimentation on a benchtop at room temperature. (B) Pictures of streptavidin-modified JAAZ (Strep-JAAZ) sample at 0, 1, 2, 4, 6, 8 and 24 hours of sedimentation on a benchtop at room temperature. Both samples are filtered and at an equivalent free dye concentration of 200 $\mu$M. The volumes of both samples are 50 $\mu$L. (n=1).

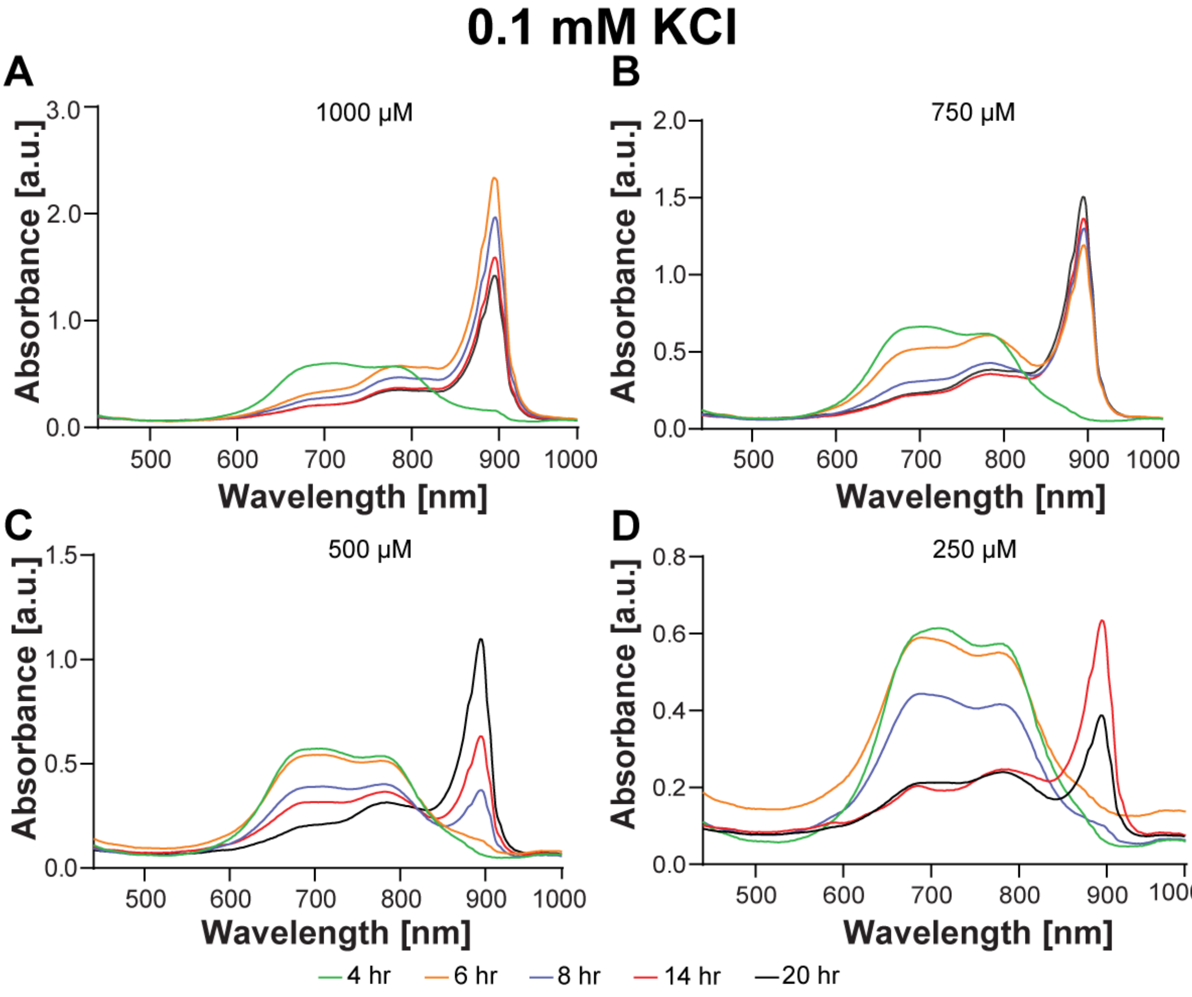


**Figure S14. Formation of JAAZ in 0.1 mM KCl at different ICG dye concentrations**. (A-D) UV-Vis-NIR spectrums of 1:10 ICG azide:ICG molar solution in 0.1 mM KCl concentration at 1000 μM A), 750 μM B), 500 μM C) and 250 μM D) total dye concentrations depicted for 20 hours.

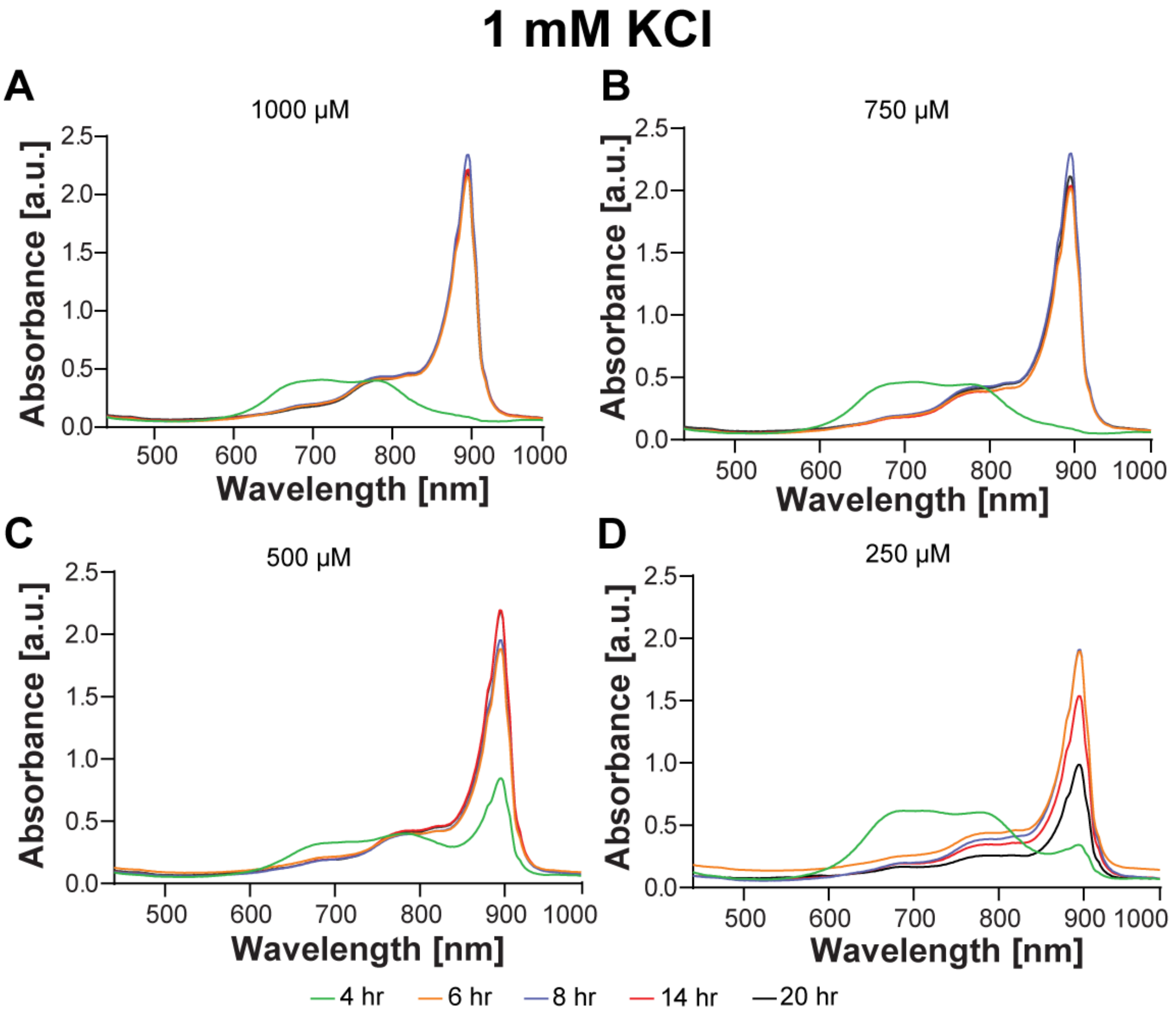


**Figure S15. Formation of JAAZ in 1 mM KCl at different ICG dye concentrations**. (A-D) UV-Vis-NIR spectrums of 1:10 ICG azide:ICG molar solution in 0.1 mM KCl concentration at 1000 $\mu$M A), 750 $\mu$M B), 500 $\mu$M C) and 250 $\mu$M D) total dye concentrations depicted for 20 hours.

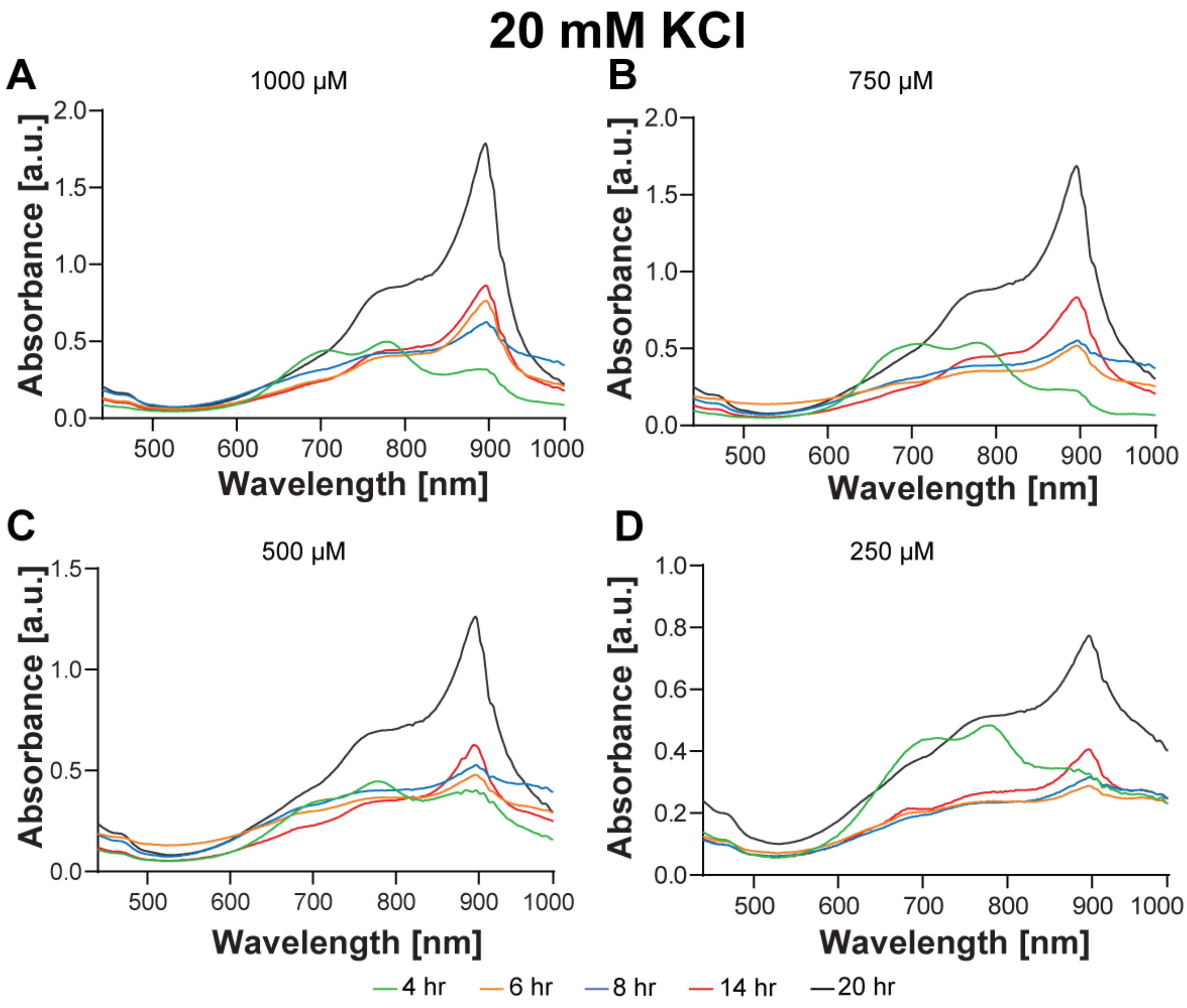


**Figure S16. Formation of JAAZ in 20 mM KCl at different ICG dye concentrations**. (A-D) UV-Vis-NIR spectrums of 1:10 ICG-azide:ICG molar solution in 20 mM KCl concentration at 1000 μM (A), 750 μM (B), 500 μM (C) and 250 μM (D) total dye concentrations depicted for 20 hours.

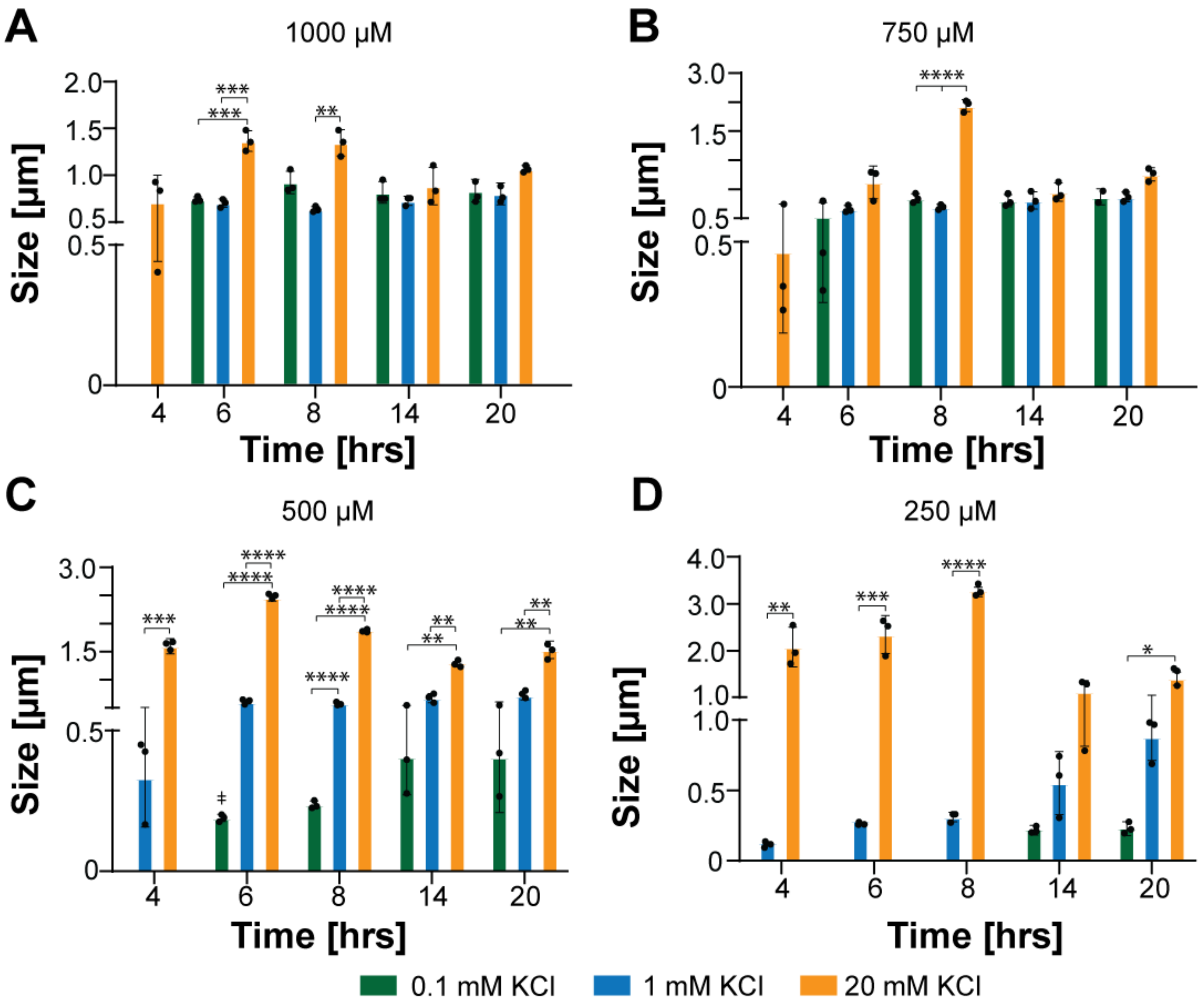


**Figure S17. Size of JAAZ in different KCl concentrations.** (A-D) Average size formed JAAZ of 1:10 molar solutions at 1000 $\mu$M (A), 750 $\mu$M (B), 500 $\mu$M (C) and 250 $\mu$M (D) total dye concentrations depicted for 20 hours. (data presented as mean±SD, n=3, p-values are calculated using one-way ANOVA, *=p<0.05, **=p<0.01, ***=p<0.001, ****=p<0.0001; all P values are provided in Table S9).

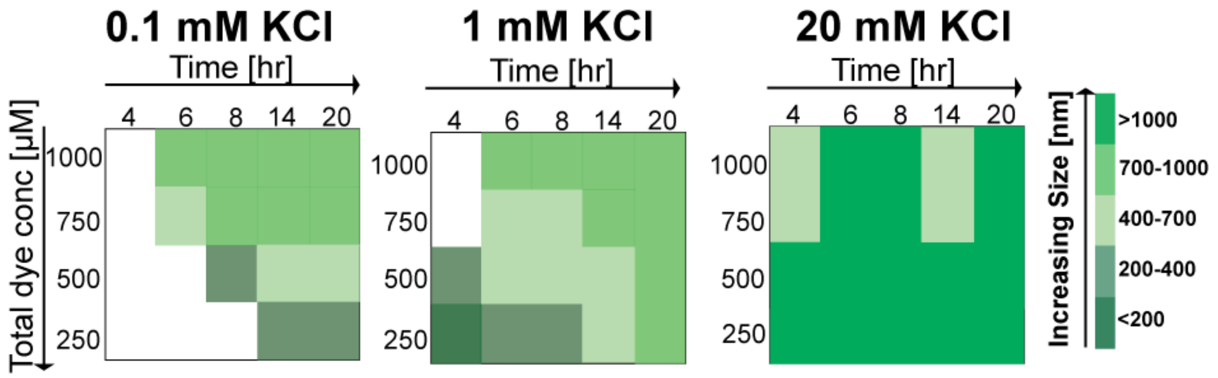


**Figure S18. Summary of effects of KCl concentration, dye concentration, and time on the average size of JAAZ particles.** The summary represents the range of sizes of JAAZ particles formed at different KCl concentrations with increasing time and increasing total dye concentration. A color bar representing the different sizes range (in nm) is also given.

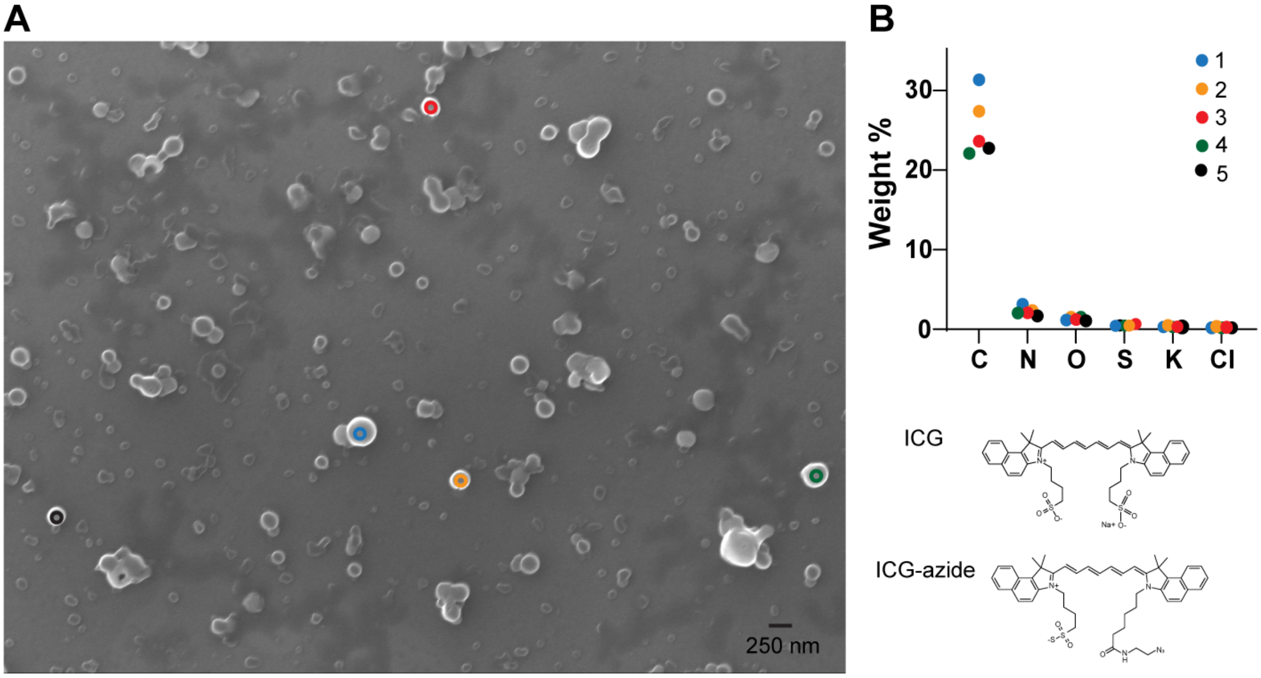


**Figure S19. SEM image and EDX analysis of N-JAAZ particles.** (A) SEM image of N-JAAZ particles prepared at 250 $\mu$M dye concentration and 0.1 mM KCl. Size measurements of N-JAAZ particles using DLS showed an average of 183 nm. (B) EDX analysis showing weight percentage of carbon (C), nitrogen (N), oxygen (O), sulfur (S), potassium (K) and chloride (Cl) of five different N-JAAZ particles as marked in image a. Chemical structures of ICG and ICG-azide dye are shown for reference.

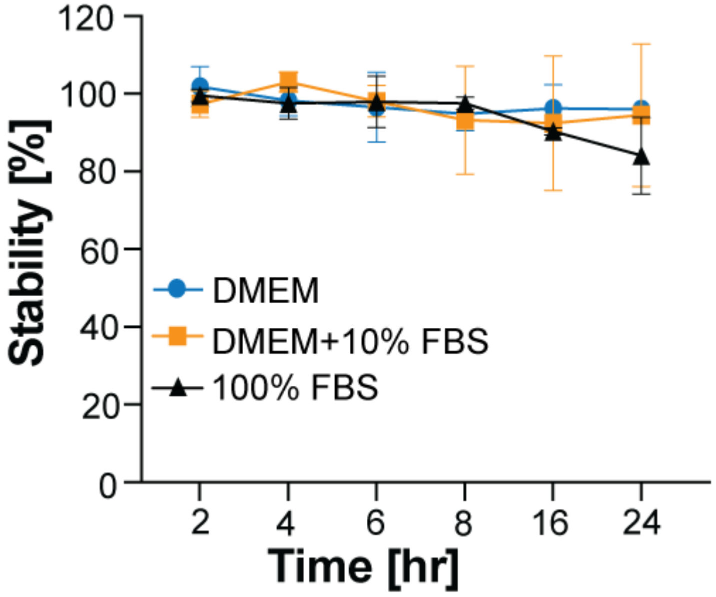


**Figure S20. Percent stability of N-JAAZ particles in media and serum.** Stability of N-JAAZ particles incubated in DMEM (blue), DMEM+10% FBS (orange) and 100% FBS (black) at 37 °C. The magnitude of the 895 nm absorbance peak was used to determine the stability of the particles. N-JAAZ particles showed ~20% decrease in the 895 nm peak at 24 hours of incubation in 100% FBS. (data represented as mean±SD, n=3 distinct replicates).

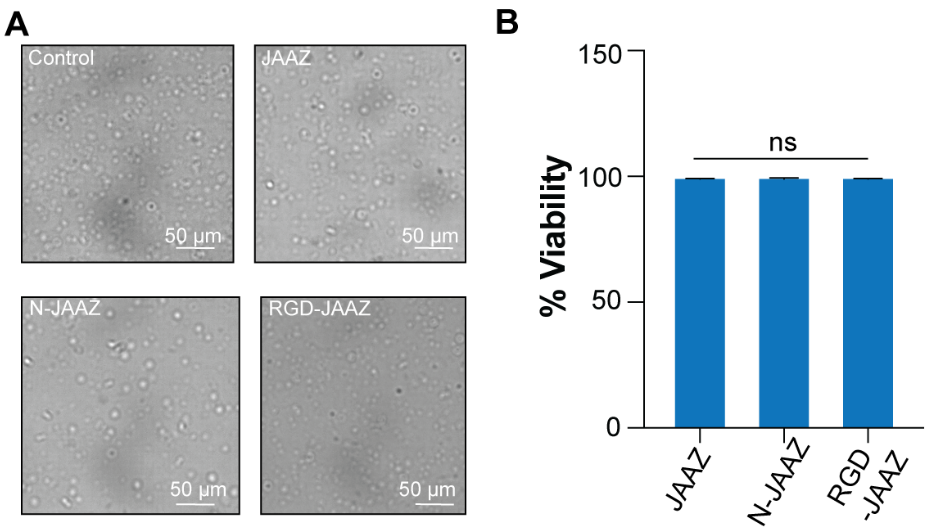

**Figure S21. Hemotoxicity and cytotoxicity of JAAZ particles.** (A) Optical microscopy images of red blood cells (RBCs) incubated with JAAZ (930±27 nm), N-JAAZ (256±21 nm) and RGD-JAAZ (997±48 nm) particles. There were no morphological changes observed of the RBCs between the control, untreated red blood cells and the treated cells. (B) MTT assay results of HeLa cells incubated with 20 $\mu$M equivalent dye concentration JAAZ, N-JAAZ and RGD-JAAZ particles (data presented as mean±SD, n=3, p-values are calculated using one-way ANOVA, ns=p=0.9531).

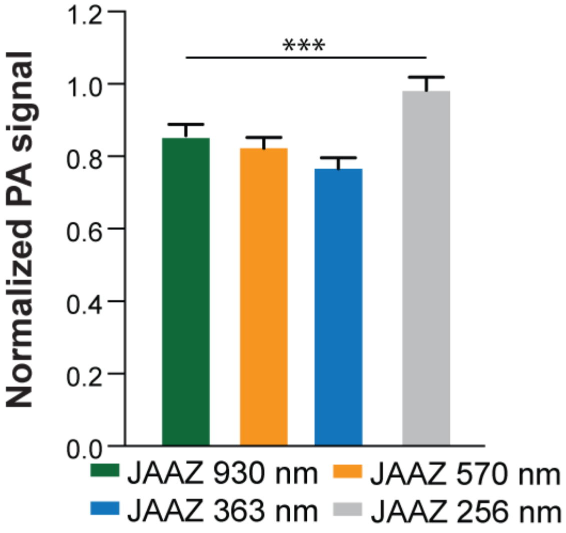


**Figure S22. PA signal amplitude of different sized JAAZ particles.** PA signal of JAAZ particles of different sizes along with N-JAAZ particles embedded in 0.5% (w/v) agarose. The sizes of the compared particles are 256±20 nm, 363±90 nm, 570±52 nm and 930±27 nm. The PA signal is normalized with respect to ICG-JA particles. All samples are at 2.0 absorbance OD. (data presented as mean±SD, n=100, p-values are calculated using one-way ANOVA, ***=p<0.001).

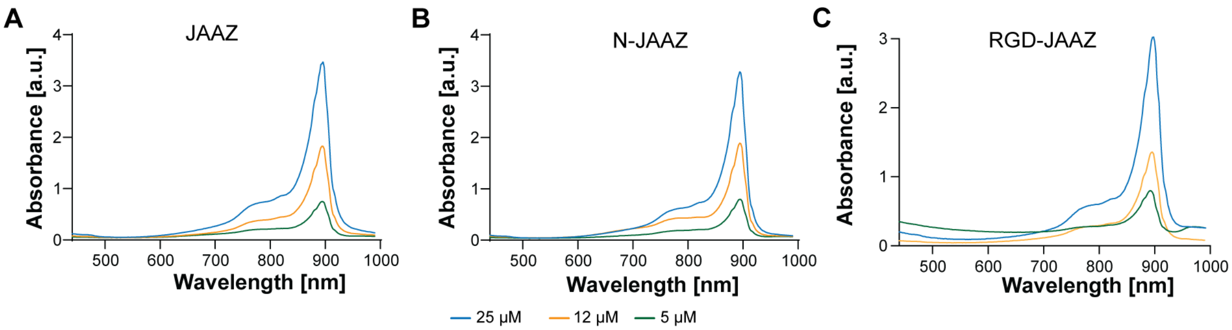


**Figure S23. UV-Vis-NIR spectra of different JAAZ samples**. (A) Absorbance spectrum of JAAZ particles at 25, 12 and 5 $\mu$M equivalent free dye concentrations. (B) Absorbance spectrum of nanoscale JAAZ (N-JAAZ) at 25, 12 and 5 $\mu$M equivalent free dye concentrations. (C) Absorbance spectrum of peptide modified JAAZ (RGD-JAAZ) at 25, 12 and 5 $\mu$M of equivalent free dye concentrations. Absorbance reading of all samples is taken at 120 $\mu$l (n=3 distinct replicates).

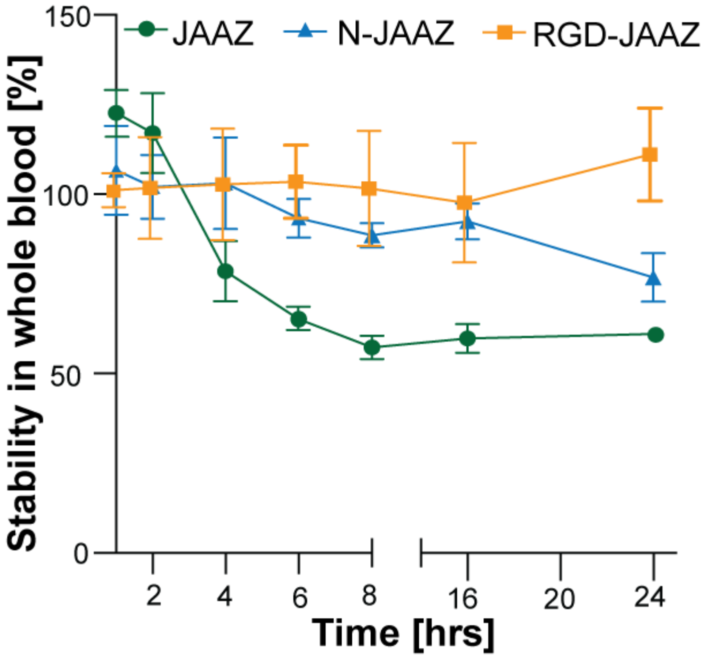


**Figure S24. Percentage stability of JAAZ particles in whole blood.** JAAZ, N-JAAZ and RGD-JAAZ particles at 10 $\mu$M equivalent free dye were incubated in whole sheep's blood at 37 °C for 24 hours. The magnitude of the 895 nm absorbance peak was used to determine the stability of the particles. RGD-JAAZ particles remained stable for 24 hours whereas N-JAAZ particles observed ~20% degradation at 24 hours of incubation. JAAZ particles showed ~30% decrease in the 895 nm peak 4 hours after incubation and were 60% stable after 24 hours. (data represented as mean±SD, n=3 distinct replicates).

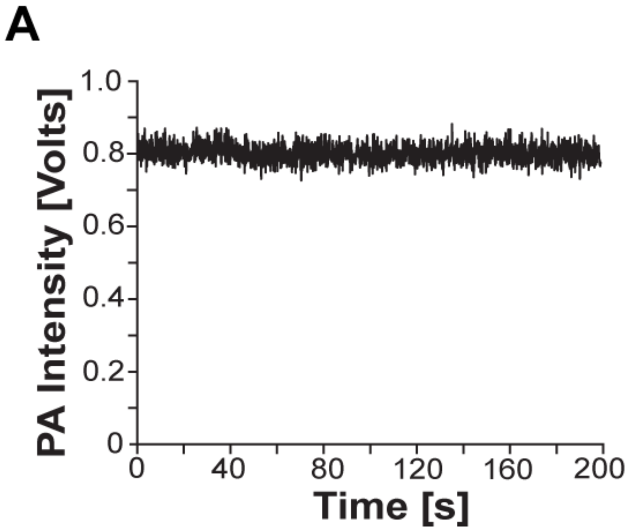


**Figure S25. Photobleaching and scan of a pattern using RGD-JAAZ**. (A) PA signal intensity of 10 μM RGD-JAAZ particles under maximum laser power. (B) 2D photoacoustic image of 'M' pattern made with 10 μM RGD-JAAZ particles.

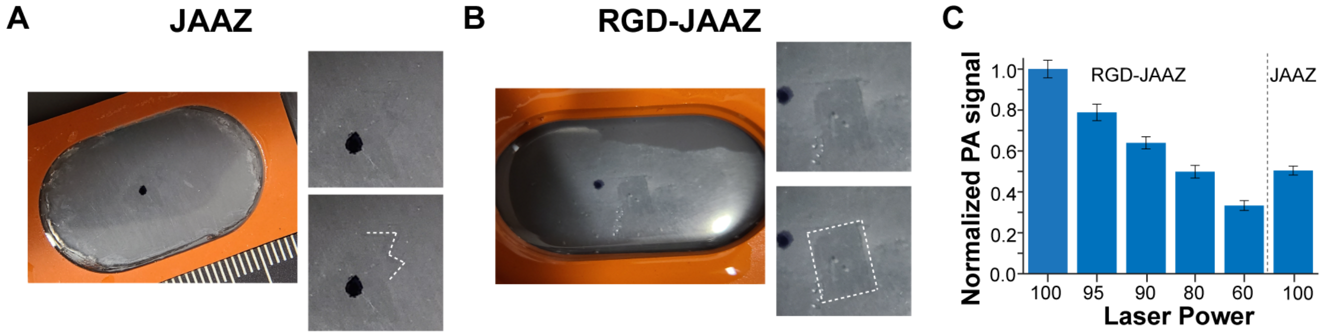


**Figure S26. Picture of HeLa cells stained using JAAZ and RGD-JAAZ and their intensity** (A) Image of a glass slide with a red silicone insert creating a well containing HeLa cells stained with 10 μM JAAZ particles. Before staining, the cells were scraped using a razor blade as indicated by white dashed lines 'M' shape overlay on the bottom inset that highlight the scrapped region. (B) Image of a glass slide with a red silicone insert creating a well containing HeLa cells stained with 10 $\mu$M RGD-JAAZ particles. Before staining, the cells were scraped using a razor blade as indicated by the white dashed lines square shape overlay on the botom inset that highlight the scrapped region. A black sharpie dot was placed on the glass slide in both (A) and (B) to help identify the postition of the pattern under the transducer while scanning. (C) Maximum PA signal from a 100 $\mu$m area of stained cells from both JAAZ and RGD-JAAZ stained glass slides at varying laser powers (100, 95, 90, 80, and 60). The PA signal is normalized with respect to the signal from RGD-JAAZ at maximum laser power.

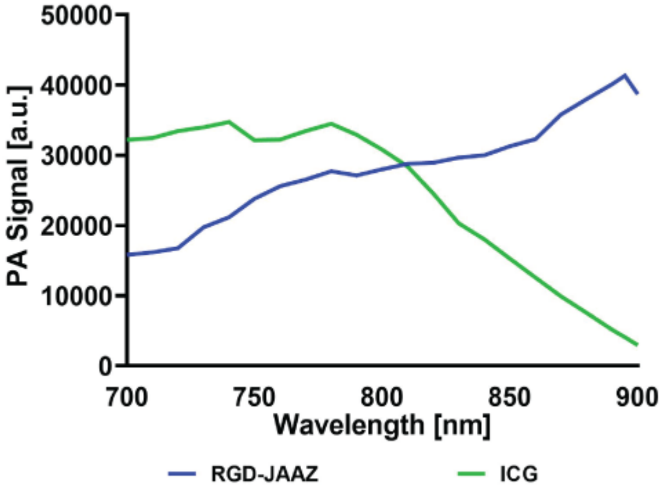


**Figure S27. Phantom imaging of RGD-JAAZ particles.** Measured PA spectra of 0.4 mM RGD-JAAZ particles and monomeric ICG dye acquired with the TriTom™ imaging system.

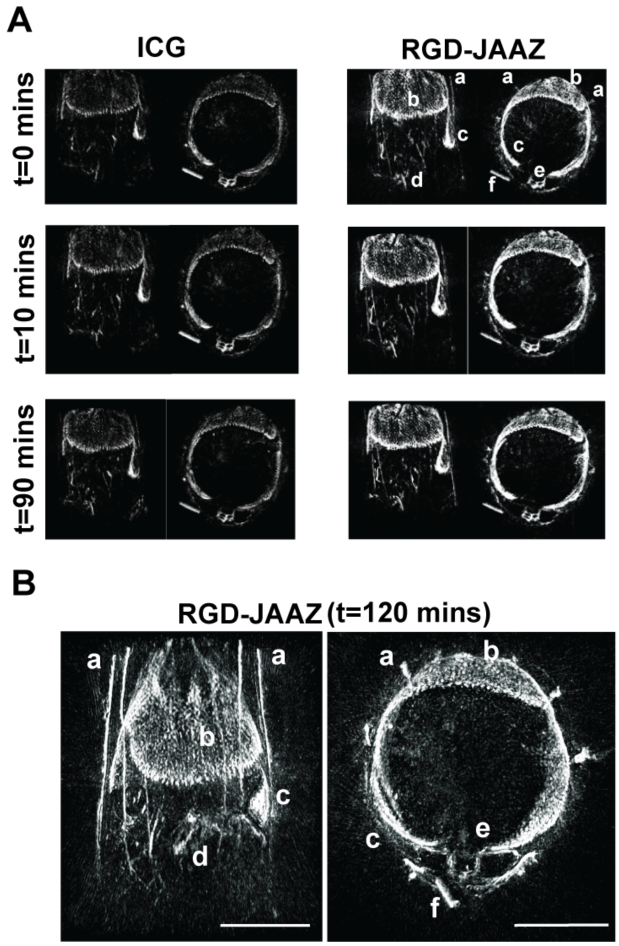


**Figure S28.** ***In vivo* imaging using RGD-JAAZ particles.** (A) Coronal and axial slabs of the ICG (left) and RGD-JAAZ particle (right) injections acquired at the peak excitation wavelength (i.e., 800 and 895 nm, respectively) over time. The displayed dynamic range for all images was set based on the baseline scan acquired at the same wavelength. (B) Representative coronal and axial slab of the RGD-JAAZ particle injection acquired at the peak excitation wavelength of 895 nm at 120 minutes. (Legend for both A and B): a. superficial thoracic vessels, b. liver c. spleen d. intestines e. thoracic vertebrae f. $CuSO_4$ fiducial tube. Scale bar for B is 10 mm.

Table S1.

Size and charge (zeta potential) of JAAZ and N-JAAZ before and after vacuum drying and resuspension in water.

| Sample | Size (nm) | | Zeta Potential (mV) | |
|---|---|---|---|---|
| | Before Drying | After Drying | Before Drying | After Drying |
| JAAZ | 1,083 ± 52 | 1,030 ± 256 | -52.1 ± 2 | -55.0 ± 5 |
| Nanosized JAAZ | 281.8 ± 37 | 360.9 ± 38 | -54.7± 1 | -57.1± 3 |

Sample size is n=3 for DLS measurement; p=0.8470=ns for JAAZ; p=0.4622=ns for N-JAAZ (Statistics are for DLS measurement); n=2 for zeta potential measurement.

Table S2.

Samples and their attached functional groups/biomolecules

| Sample Name | Composition and Functional Groups Attached | | | | |
|---|---|---|---|---|---|
| | ICG (mM) | $ICG-N_3$ (mM) | DBCO-Biotin (Molar ratio with respect to $ICG-N_3$) | Streptavidin | RGD-Biotin |
| JA | 1 | - | - | - | - |
| JAAZ | 1 | 0.1 | - | - | - |
| Bio-JAAZ | 1 | 0.1 | 10X | - | - |
| Strep-JAAZ | 1 | 0.1 | 10X | 10X | - |
| RGD-JAAZ | 1 | 0.1 | 10X | 10X | 10X |

Table S3.

P-values for Fig. 2.

| Fig 2A | | |
|---|---|---|
| Row | JA | JAAZ |
| JA | Na | 0.0001 |
| JAAZ | 0.0001 | Na |

| Fig 2B | | |
|---|---|---|
| Row | JA | JAAZ |
| JA | Na | 0.0161 |
| JAAZ | 0.0161 | Na |

Table S4.

P-values for Fig. 3.

| Fig. 3C | | | | |
|---|---|---|---|---|
| Row | JAAZ | Bio-JAAZ | Strep-JAAZ | RGD-JAAZ |
| JAAZ | Na | 0.9058 | 0.0013 | 0.0019 |
| Bio-JAAZ | 0.9058 | Na | <0.0001 | <0.0001 |
| Strep-JAAZ | 0.0013 | <0.0001 | Na | 0.9011 |
| RGD-JAAZ | 0.0019 | <0.0001 | 0.9011 | Na |

Table S5.

P-values for Fig. 4

| Fig. 4B | | | |
|---|---|---|---|
| Row | 0.1 mM KCl | 1 mM KCl | 20 mM KCl |
| 0.1 mM KCl | Na | 0.0007 | <0.0001 |
| 1 mM KCl | 0.0007 | Na | <0.0001 |
| 20 mM KCl | <0.0001 | <0.0001 | Na |

| Fig. 4E | | | | |
|---|---|---|---|---|
| Row | N-JAAZ | N-Bio-JAAZ | N-Strep-JAAZ | N-RGD-JAAZ |
| N-JAAZ | Na | 0.3627 | 0.3634 | 0.0690 |
| N-Bio-JAAZ | 0.3627 | Na | 0.0645 | 0.0882 |
| N-Strep-JAAZ | 0.3634 | 0.0645 | Na | 0.0919 |
| N-RGD-JAAZ | 0.0690 | 0.0882 | 0.0919 | Na |

| Fig. 4F | | | | |
|---|---|---|---|---|
| Row | N-JAAZ | N-Bio-JAAZ | N-Strep-JAAZ | N-RGD-JAAZ |
| N-JAAZ | Na | 0.7620 | 0.0007 | 0.0004 |
| N-Bio-JAAZ | 0.7620 | Na | 0.0004 | 0.0002 |
| N-Strep-JAAZ | 0.0007 | 0.0004 | Na | 0.4006 |
| N-RGD-JAAZ | 0.0004 | 0.0002 | 0.4006 | Na |

.

Table S6.

P-values for Fig. 5

| Fig. 5B, 25 $\mu$M | | |
|---|---|---|
| Row | JAAZ | N-JAAZ |
| JAAZ | Na | <0.0001 |
| N-JAAZ | <0.0001 | Na |

| Fig. 5B, 12 $\mu$M | | | |
|---|---|---|---|
| Row | JAAZ | N-JAAZ | RGD-JAAZ |
| JAAZ | Na | <0.0001 | <0.0001 |
| N-JAAZ | <0.0001 | Na | <0.0001 |
| RGD-JAAZ | <0.0001 | <0.0001 | Na |

| Fig. 5B, 5 $\mu$M | | | |
|---|---|---|---|
| Row | JAAZ | N-JAAZ | RGD-JAAZ |
| JAAZ | Na | <0.0001 | <0.0001 |
| N-JAAZ | <0.0001 | Na | <0.0001 |
| RGD-JAAZ | <0.0001 | <0.0001 | <0.0001 |

| Fig. 5D, 25 $\mu$M | | | | |
|---|---|---|---|---|
| Row | ICG | JAAZ | N-JAAZ | RGD-JAAZ |
| ICG | Na | <0.0001 | <0.0001 | <0.0001 |
| JAAZ | <0.0001 | Na | <0.0001 | <0.0001 |
| N-JAAZ | <0.0001 | <0.0001 | Na | <0.0001 |
| RGD-JAAZ | <0.0001 | <0.0001 | <0.0001 | Na |

Table S7.

P-values for Fig. S11

| Fig. S11A | | | | |
|---|---|---|---|---|
| Row | JAAZ | Bio-JAAZ | Strep-JAAZ | RGD-JAAZ |
| JAAZ | Na | 0.5494 | 0.8409 | 0.0940 |
| Bio-JAAZ | 0.5494 | Na | 0.5175 | 0.1036 |
| Strep-JAAZ | 0.8409 | 0.5175 | Na | 0.2148 |
| RGD-JAAZ | 0.0940 | 0.1036 | 0.2148 | Na |

| Fig. S11B | | | | |
|---|---|---|---|---|
| Row | JAAZ | Bio-JAAZ | Strep-JAAZ | RGD-JAAZ |
| JAAZ | Na | 0.9729 | 0.6851 | 0.3637 |
| Bio-JAAZ | 0.9729 | Na | 0.7482 | 0.4650 |
| Strep-JAAZ | 0.7482 | 0.7482 | Na | 0.7118 |
| RGD-JAAZ | 0.3637 | 0.4650 | 0.7118 | Na |

Table S8.

P-values for Fig. S12

| Fig. S12B | | | | |
|---|---|---|---|---|
| Row | 1:20 | 1:10 | 1:5 | 1:3 |
| 1:20 | Na | 0.2901 | 0.7246 | 0.2769 |
| 1:10 | 0.2901 | Na | 0.4064 | 0.7123 |

| | | | | |
|---|---|---|---|---|
| 1:5 | 0.7246 | 0.4064 | Na | 0.4275 |
| 1:3 | 0.2749 | 0.7123 | 0.4275 | Na |

Table S9.

P-values for Fig. S17

| Fig. S17, 1000 $\mu$M, 6 hrs | | | |
|---|---|---|---|
| Row | 0.1 mM | 1 mM | 20 mM |
| 0.1 mM | Na | 0.2833 | 0.0007 |
| 1 mM | 0.2833 | Na | 0.0006 |
| 20 mM | 0.0007 | 0.0006 | Na |

| Fig. S17, 1000 $\mu$M, 8 hrs | | | |
|---|---|---|---|
| Row | 0.1 mM | 1 mM | 20 mM |
| 0.1 mM | Na | 0.0157 | 0.0165 |
| 1 mM | 0.0157 | Na | 0.0010 |
| 20 mM | 0.0165 | 0.0010 | Na |

| Fig. S17, 1000 $\mu$M, 14 hrs | | | |
|---|---|---|---|
| Row | 0.1 mM | 1 mM | 20 mM |
| 0.1 mM | Na | 0.2834 | 0.6409 |
| 1 mM | 0.2834 | Na | 0.2570 |
| 20 mM | 0.6409 | 0.2570 | Na |

| Fig. S17, 1000 $\mu$M, 20 hrs | | | |
|---|---|---|---|
| Row | 0.1 mM | 1 mM | 20 mM |
| 0.1 mM | Na | 0.7945 | 0.0462 |

| | | | |
|---|---|---|---|
| 1 mM | 0.7945 | Na | 0.0296 |
| 20 mM | 0.0462 | 0.0296 | Na |

| Fig. S17, 750 $\mu$M, 6 hrs | | | |
|---|---|---|---|
| Row | 0.1 mM | 1 mM | 20 mM |
| 0.1 mM | Na | 0.4137 | 0.0506 |
| 1 mM | 0.4137 | Na | 0.0502 |
| 20 mM | 0.0506 | 0.0502 | Na |

| Fig. S17, 750 $\mu$M, 8 hrs | | | |
|---|---|---|---|
| Row | 0.1 mM | 1 mM | 20 mM |
| 0.1 mM | Na | 0.0428 | 0.00001 |
| 1 mM | 0.0428 | Na | 0.0502 |
| 20 mM | 0.00002 | 0.00001 | Na |

| Fig. S17, 750 $\mu$M, 14 hrs | | | |
|---|---|---|---|
| Row | 0.1 mM | 1 mM | 20 mM |
| 0.1 mM | Na | 0.9776 | 0.2496 |
| 1 mM | 0.0428 | Na | 0.3148 |
| 20 mM | 0.2496 | 0.3148 | Na |
| Fig. S17, 750 $\mu$M, 20 hrs | | | |
| Row | 0.1 mM | 1 mM | 20 mM |
| 0.1 mM | Na | 0.9776 | 0.0421 |
| 1 mM | 0.9776 | Na | 0.0089 |
| 20 mM | 0.0421 | 0.0089 | Na |

| Fig. S17, 500 $\mu$M, 4 hrs | | |
|---|---|---|
| Row | 1 mM | 20 mM |
| 1 mM | Na | 0.0005 |
| 20 mM | 0.0005 | Na |

| Fig. S17, 500 $\mu$M, 6 hrs | | | |
|---|---|---|---|
| Row | 0.1 mM | 1 mM | 20 mM |
| 0.1 mM | Na | 0.00003 | 0.00007 |
| 1 mM | 0.00003 | Na | 0.00002 |
| 20 mM | 0.00007 | 0.00002 | Na |

| Fig. S17, 500 $\mu$M, 8 hrs | | | |
|---|---|---|---|
| Row | 0.1 mM | 1 mM | 20 mM |
| 0.1 mM | Na | 0.00007 | <0.00001 |
| 1 mM | 0.00007 | Na | 0.00002 |
| 20 mM | <0.00001 | 0.00002 | Na |

| Fig. S17, 500 $\mu$M, 14 hrs | | | |
|---|---|---|---|
| Row | 0.1 mM | 1 mM | 20 mM |
| 0.1 mM | Na | 0.0419 | 0.0030 |
| 1 mM | 0.0419 | Na | 0.0022 |
| 20 mM | 0.0030 | 0.0022 | Na |

| Fig. S17, 500 $\mu$M, 20 hrs | | | |
|---|---|---|---|
| Row | 0.1 mM | 1 mM | 20 mM |
| 0.1 mM | Na | 0.0785 | 0.0055 |

| | | | |
|---|---|---|---|
| 1 mM | 0.0785 | Na | 0.0013 |
| 20 mM | 0.0055 | 0.0013 | Na |

| Fig. S17, 500 $\mu$M, 4 hrs | | |
|---|---|---|
| Row | 1 mM | 20 mM |
| 1 mM | Na | 0.0013 |
| 20 mM | 0.0013 | Na |

| Fig. S17, 500 $\mu$M, 6 hrs | | |
|---|---|---|
| Row | 1 mM | 20 mM |
| 1 mM | Na | 0.0009 |
| 20 mM | 0.0009 | Na |

| Fig. S17, 500 $\mu$M, 8 hrs | | |
|---|---|---|
| Row | 1 mM | 20 mM |
| 1 mM | Na | 0.00003 |
| 20 mM | 0.00003 | Na |

| Fig. S17, 500 $\mu$M, 14 hrs | | | |
|---|---|---|---|
| Row | 0.1 mM | 1 mM | 20 mM |
| 0.1 mM | Na | 0.1494 | 0.0322 |
| 1 mM | 0.1494 | Na | 0.0627 |
| 20 mM | 0.0322 | 0.0627 | Na |

| Fig. S17, 500 $\mu$M, 20 hrs | | | |
|---|---|---|---|
| Row | 0.1 mM | 1 mM | 20 mM |

| | | | |
|---|---|---|---|
| 0.1 mM | Na | 0.0149 | 0.0186 |
| 1 mM | 0.0149 | Na | 0.0530 |
| 20 mM | 0.0186 | 0.0530 | Na |
